# Small RNAs in plasma extracellular vesicles define biomarkers of premanifest changes in Huntington’s disease

**DOI:** 10.1101/2023.12.01.568823

**Authors:** Marina Herrero-Lorenzo, Jesús Pérez-Pérez, Georgia Escaramís, Saül Martínez-Horta, Rocío Pérez-González, Elisa Rivas-Asensio, Jaime Kulisevsky, Ana Gámez-Valero, Eulàlia Martí

## Abstract

Despite the advances in the understanding of Huntington’s disease (HD), there is the need for molecular biomarkers to categorize mutation-carriers during the preclinical stage of the disease preceding the functional decline. Small RNAs (sRNAs) are a promising source of biomarkers since their expression levels are highly sensitive to pathobiological processes. Here, using an optimized method for plasma extracellular vesicles (EVs) purification and an exhaustive analysis pipeline of sRNA sequencing data, we show that EV-sRNAs are early downregulated in mutation-carriers, and that this deregulation is associated with premanifest cognitive performance. Seven candidate sRNAs (tRF-Glu-CTC, tRF-Gly-GCC, miR-451a, miR-21-5p, miR-26a-5p, miR-27a-3p, and let7a-5p) were validated in additional subjects, showing a significant diagnostic accuracy at premanifest stages. Of these, miR – 21-5p was significantly decreased over time in a longitudinal study; and miR-21-5p and miR-26a-5p levels correlated with cognitive changes in the premanifest cohort. In summary, the present results suggest that deregulated plasma EV-sRNAs define an early biosignature in mutation carriers with specific species sensing the progression and cognitive changes occurring at the premanifest stage.

## Introduction

Huntington’s disease (HD) is a dominantly inherited neurodegenerative disorder caused by a CAG triplet repeat expansion in the Huntingtin gene (*HTT*)^1^. The main neuropathological hallmarks of HD are the presence of cytoplasmatic mutant HTT aggregates, the massive loss of the medium spiny neurons of the striatum, and cortical degeneration^2^. In current clinical practice, manifest HD (M-HD) patients are diagnosed based on the appearance of a complex constellation of clinical symptoms, including progressive motor abnormalities, neuropsychiatric disturbances, early cognitive deterioration, and dementia, among others^3,4^. However, genetic analysis allows the use of predictive testing to identify premanifest gene-mutation carriers (P-HD) who may exhibit progressive brain changes in specific imaging metrics and cognitive performance long before the estimated time to diagnosis^5,6^.

Current treatments for HD provide only symptomatic relief, and later results on clinical trials investigating disease-modifying therapies have not shown significant efficacy so far ^7^. The combination of predictive genetic testing with new early molecular biomarkers could help in improving the design of clinical trials for HD by selecting appropriate patient populations, stratifying patients for intervention, monitoring treatment response, and improving trial efficiency.

Later efforts have been invested in the definition of HD biofluid-based biomarkers. The most remarkable one is mutant HTT, which can be measured in the cerebrospinal fluid (CSF) to assess the efficacy of mutant HTT-lowering therapies^8^. Further, it has been shown that minimally invasive plasma biomarkers can reflect molecular changes occurring in this heterogeneous disorder. Specifically, plasma neurofilament light chain (NfL) shows a higher prognostic value than mutant HTT^9,10^, despite not being disease-specific, as it is also observed in other brain-related diseases^11,12^. Otherwise, regarding RNA-based biomarkers, microRNAs (miRNAs) have been largely studied as promising transcriptomic biomarkers since they are stable in peripheral circulation^13^, and their levels are highly sensitive to physiological and pathological states^14–16^. Moreover, we have recently shown that not only miRNAs but other small RNAs (sRNAs, < 200 nucleotides) may have a profound role in HD pathogenesis^17^. The intrastriatal injection of sRNAs from HD patients’ brains triggers motor abnormalities, neuronal toxicity, and transcriptional alterations in naïve mice^17^. These results suggest that different types of sRNAs are sufficient to induce neurotoxicity and are likely involved in the progressive transcriptomic deregulation underlying HD pathogenesis. Therefore, a better understanding of sRNA expression dynamics should provide clues about the progression of the disease and help to elucidate if sRNA perturbations are an early phenomenon in HD.

In peripheral circulation, extracellular sRNAs (exRNAs) can be found freely circulating, associated to lipid-or protein-complexes, or encapsulated within different types of vesicles. Specifically, extracellular vesicles (EVs) are bilayered vesicles secreted by many cell types that encapsulate molecules such as RNAs and proteins from the cell of origin, which makes them attractive in biomarker discovery^18–21^. Nevertheless, recent studies have raised controversy as they show that the vast majority of circulating exRNAs are not found in EVs, but co-purifying with other molecules in the extravesicular milieu^22,23^. In plasma, proteins such as Argonaute2 ribonucleic complexes have been described as important miRNA carriers^23^; however, no solid evidence regarding the distribution of sRNA biotypes in the plasma subfractions has been described so far.

Herein, we aimed to define biomarkers to monitor changes that occur in the pre-symptomatic HD population, by exploring plasma sRNA levels. We analyzed sRNA profiles of EVs-enriched and NonEV plasma fractions from P-HD and M-HD mutation carriers in comparison to control samples. Our data indicate that most exRNAs in plasma EVs are downregulated in mutation carriers, with many changes occurring at premanifest stages. Importantly, specific sRNAs showing altered expression over time correlate with premanifest cognitive performance, which may lead to the definition of novel early progression biomarkers.

## Results

### 1. Different plasma subfractions show specific sRNA profiles

Aiming to understand exRNA distribution in plasma and identify informative compartments, we profiled sRNAs associated with different plasma subfractions. To this end five plasma samples from healthy volunteers were subfractionated through SEC-UF^24^ (Fig.1A). As described previously^25,26^, SEC-eluted fractions enriched in EVs were determined by flow cytometry, and the four fractions showing positivity for tetraspanins CD9, CD63, and CD81 were pooled (Fig. 1B). Subsequently, tetraspanins-positive pooled fractions were concentrated by UF obtaining a concentrated EVs-enriched fraction (cEVs) and an eluted fraction without EVs (Filtrate). In parallel, tetraspanins-negative SEC-eluted fractions showed an increasing protein concentration (NonEVs fractions) and we selected two different grouped types of fractions: fractions with low concentration of protein (NonEVs-Low) and fractions showing the highest concentration of protein (NonEVs-High) (Fig.1A).

**Figure 1:**
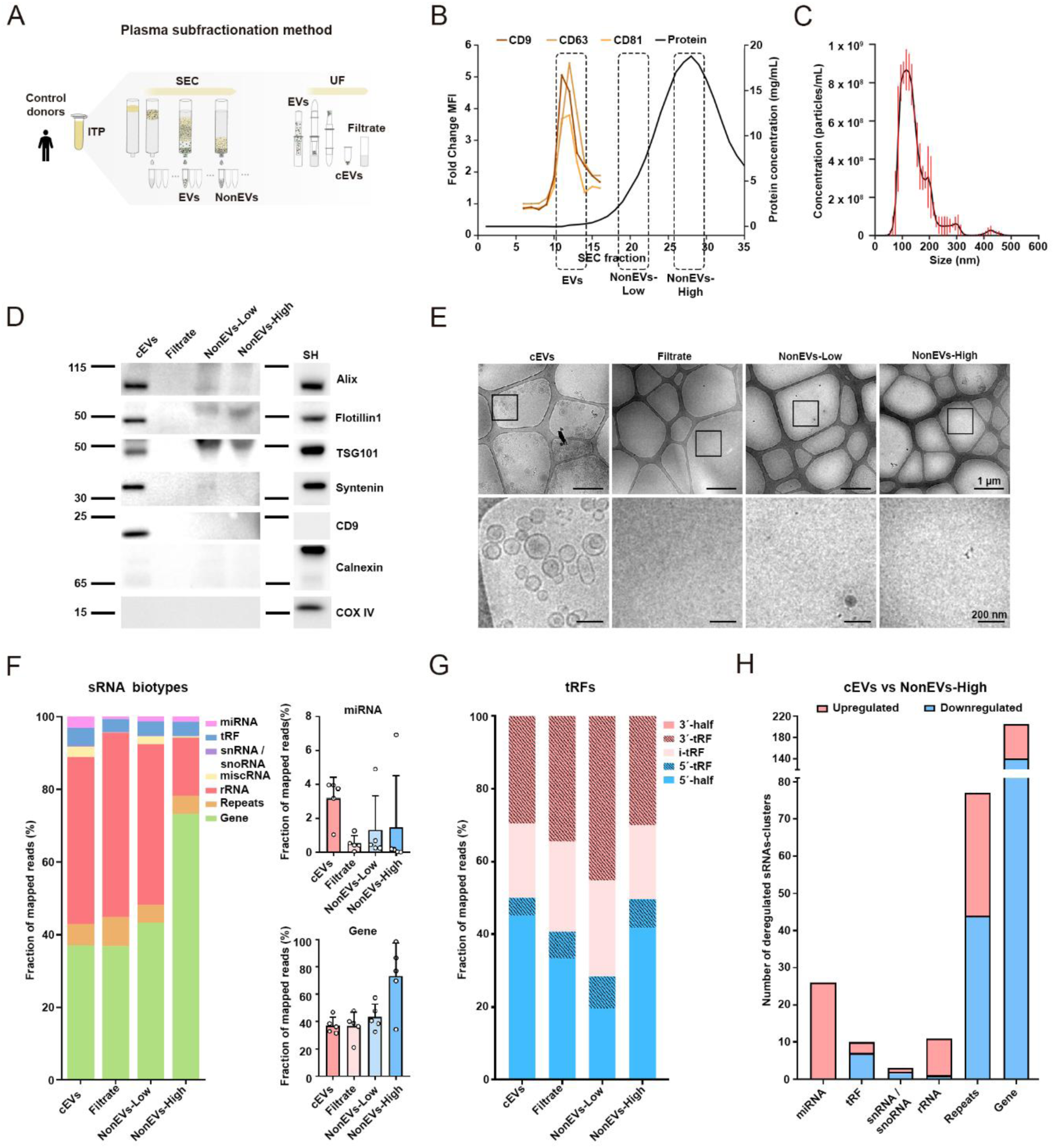
sRNA biotypes are differently distributed between plasma subfractions. (A) Schematic representation of plasma-EV isolation procedure by SEC and UF and selected fractions for analysis: cEVs, Filtrate, NonEVs-Low and NonEVs-High fractions. (B) Representative SEC profile of plasma samples showing EV elution in low-protein fractions as measured by Bradford protein assay and enriched in CD9, CD63 and CD81 markers as analyzed by flow cytometry. (C) Representative NTA size distribution profile of cEVs fraction (n=5). (D) Western Blot analysis of isolated fractions. Fractions were analyzed for the presence of EV markers (Alix, Flotillin1, TSG101, Syntenin and CD9) and negative EV-markers (Calnexin and COX IV). SH-SY5Y cell lysate (SH) was used as positive controls. (E) Representative images of cryo-EM of the isolated fractions. Upper images, bar=1 μm. Lower images, bar=200nm. Using SeqCluster tool: (F) Fraction of reads that align to small RNA types are shown per plasma subfraction. Mean ± SD is shown. (G) Classification of reads that align to tRFs in 3’-half, 3’-tRF, i-tRF, 5’-tRF, and 5’-half per plasma subfraction. (H) Total number of sRNAs-clusters differentially expressed between cEVs and NonEVs-High subfractions (p-adj<0.05). Samples of each subfraction, n=5.

cEVs were further characterized in compliance with the MISEV guidelines^27^. EV concentration yielded 8.33 x 10^9^ ± 2.65 x 10^8^ particles/mL with a mean diameter size of 146.0 ± 2.8 nm in cEVs fractions as determined by NTA (Fig. 1C). Western Blot was used to confirm the presence of additional EV markers in cEVs fractions and the absence of them in Filtrate, NonEVs-Low, and NonEVs-High fractions, as well as the lack of cell markers for intracellular contamination (Fig. 1D). The reduction of Albumin and IgGs in cEVs fractions was confirmed by Ponceau red staining of the membrane ( Supplementary data Fig. 1A). Cryo-TEM images confirmed the expected range of sizes and morphological features in cEV fractions, and absence of EVs in Filtrate, NonEVs-Low and NonEVs-High fractions (Fig. 1E). Overall, this setup indicates that we have successfully obtained differentiated plasma subfractions with the potential to show specific sRNAs profiles.

To uncover the sRNA diversity in the different plasma subfractions, we used SeqCluster^28^ and ExceRpt^29^ bioinformatic tools, offering complementary information (see Methods). We detected heterogeneous proportions of sRNAs biotypes in the different plasma fractions, dominated by rRNA– and gene fragments (Fig. 1F, Supplementary data Fig. 2A, Supplementary data Fig. 3A). We noted that cEVs fractions contained a higher percentage of reads mapping onto miRNAs compared to Filtrate, NonEVs-Low, and NonEVs-High fractions. However, the highest proportion of reads mapping onto gene fragments was found in NonEVs-High fractions (Fig. 1F). Other sRNAs biotypes, such as tRNA fragments (tRFs), were similarly abundant in the four fractions, while rRNAs and repeats fragments were found in less proportion in NonEVs-High fractions. In addition, tRF were classified based on where they map onto the precursor tRNA molecule (3’-half, 3’-tRF, i-tRF, 5’-tRF, and 5’-half)^30^ and similar distributions were observed in the different subfractions (Fig. 1G). Focusing on the total number of different sRNA biotypes identified in each subfraction, we observed that cEVs fractions contained an increased number of miRNAs (Supplementary data Fig. 2B). This result indicates that, although miRNAs were a minor proportion of the total sRNAs, they were concentrated in cEVs compared with other fractions. Despite the higher abundance of gene fragments in NonEVs-High fractions, a similar diversity was detected in the different plasma subfractions ( Supplementary data Fig. 2B, Supplementary data Fig.3C). Moreover, the increased proportion of gene fragments in NonEVs-High fraction was mainly due to protein coding and retained intron fragments (Supplementary data Fig.3B).

Through hierarchical clustering analysis, we likewise assessed the differential sRNA transcriptome in the different fractions: heatmap analysis based on miRNAs levels showed that cEVs fractions grouped in a common cluster (Supplementary data Fig. 2C, Supplementary data Fig. 3E). Similarly, 4 out of 5 NonEVs-High fractions were clustered when using the gene fragment count matrix (Supplementary data Fig. 2D, Supplementary data Fig. 3F). Of notice, differential expression analysis identified 26 upregulated and no downregulated miRNA-clusters, and 61 upregulated and 139 downregulated gene fragments-clusters in cEVs in comparison to NonEVs-High fractions (Fig. 1H, Supplementary data Fig. 3D). These findings suggest that both cEVs and NonEVs-High fractions show distinct patterns of sRNA and may provide complementary information.

### 2. No major differences were found in plasma EV morphology and concentration between CTL and *HTT* mutation carriers

Once the general profiling of sRNA diversity in plasma subfractions was established, we next sought to analyze the expression pattern of plasma exRNAs in *HTT* mutation carriers. For this purpose, three different groups of samples were considered (n=10 samples per group): P-HD, M-HD, and healthy control volunteers (CTL) (Table 1). We then selected two well-differentiated plasma fractions, the cEVs enriched fraction (EV), and a fraction depleted of cEVs and showing a high protein concentration (NonEV) (Fig. 2A). EVs were further characterized based on MISEV guidelines^27^ and compared between the three groups. Regarding tetraspanins expression, we did not observe differences between groups (Fig. 2B, 2C). Western Blot was used to confirm the presence of TSG101, Flotillin1, and Alix, and the absence of Calnexin and COX IV in representative EVs (Fig. 2D). Moreover, Ponceau red staining in the two fractions suggested strong reduction of highly abundant Albumin and IgGs in cEV fractions (Supplementary data Fig. 4A). Likewise, no statistical differences were observed in EVs concentration between groups as measured by NTA (Fig. 2E); although a trend for an increased concentration in the M-HD group is observed, in line with an analogous trend in the tetraspanins levels (Fig. 2C). The analysis of NTA concentrations divided by discrete sizes revealed significant differences in the size distribution between P-HD and M-HD in comparison to CTL samples. Specifically, an increase in vesicles ranging 80-120 nm and a decrease in vesicles >200 nm is shown (Supplementary data Fig. 4B). Using cryo-EM, we confirmed the integrity of EVs (Fig. 2F, Supplementary data Fig. 4C); however, we could not validate differences between groups at any size range (Fig. 2G). In NonEVs fractions, the measurement of total protein concentration in the three groups rendered similar results (Fig. 2H).

**Figure 2:**
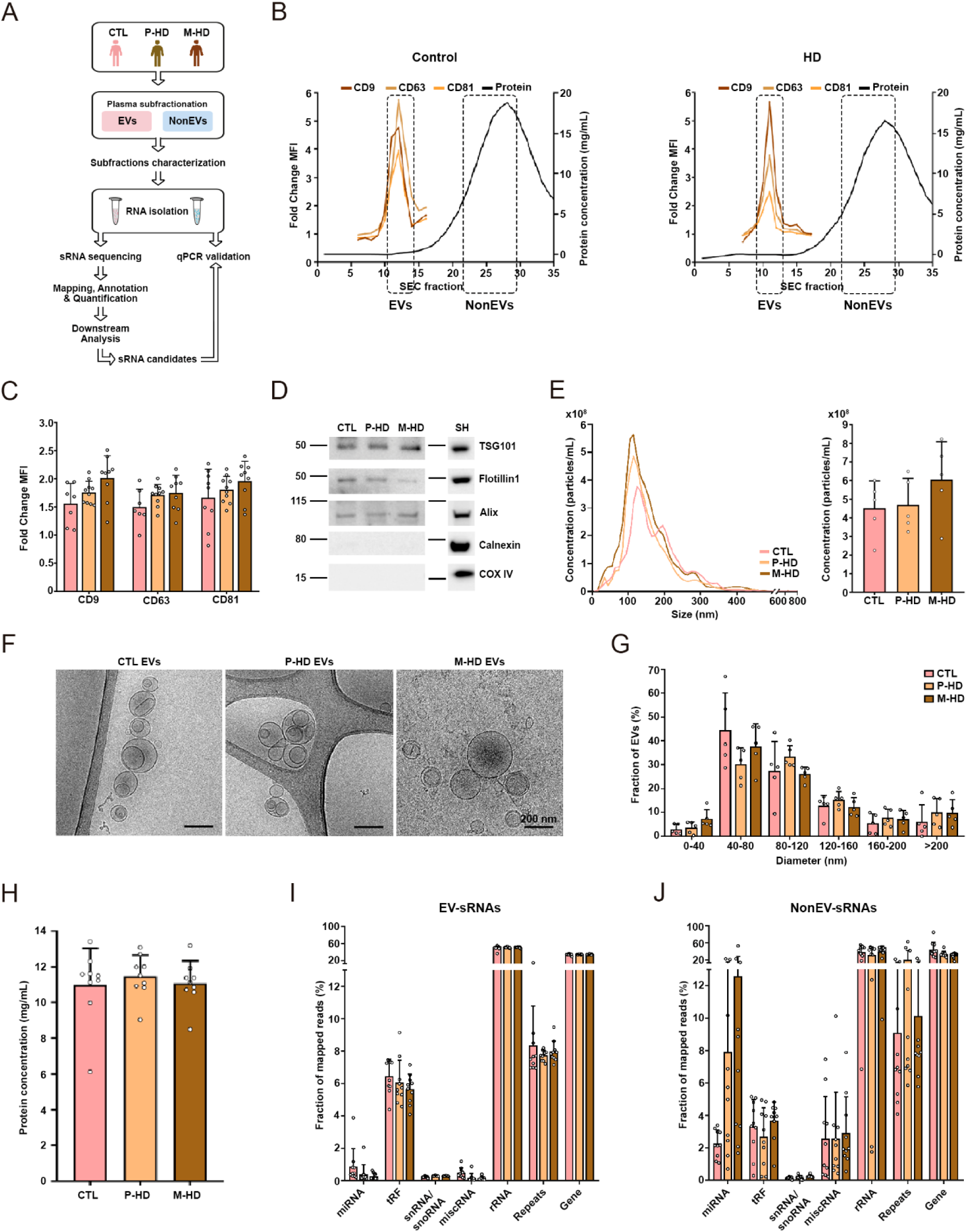
Characterization of EV fraction and NonEV fraction isolated from P-HD, M-HD, and CTL plasma. (A) Schematic representation of the workflow followed in both plasma subfractions. (B) Representative profile of CTL and HD plasma SEC fractions by protein determination and EVs fractions eluted in low protein fractions were detected using EV-markers CD9, CD63 and CD81 by flow cytometry analysis. (C) Fold Change of MFI values (MFI values compared to the isotype control – IgG-) for CD9, CD63 and CD81 in EVs pooled fractions (n=8-10 per group). (D) Representative western Blot analysis of EVs fractions for the presence of EV markers (Flotillin1, TSG101 and Alix) and negative EV-markers (COX IV and Calnexin). SH-SY5Y cell lysate (SH) was used as positive control (n=3 per group). (E) NTA size distribution profiles of EVs fractions (left) and quantification of EV particle concentration (right) (CTL vs P-HD, adj P=0.984; CTL vs M-HD, adj P=0.348; P-HD vs M-HD, adj P=0.432; n=5 per group). (F) Representative images of cryo-EM of P-HD, M-HD, and CTL EVs. Bar=200 nm (n=5 per group). (G) Size distribution of EVs diameter by cryo-EM (n=5 per group). (H) Protein content of NonEVs fractions (n=8-10 per group). sRNA profiles analyzed by SeqCluster tool showing the fraction of reads that align to small RNA types per plasma group (I) in EVs fractions and (J) NonEVs fractions (n=10 per group). Coefficient of variance (CV) in EV fraction=0.09; CV in NonEVs fraction=1.1; Levenés test used to assess the equality of variances, between EVs and NonEVs (p-val=0.0027). All data are represented as Mean ± SD.

**Table 1:**
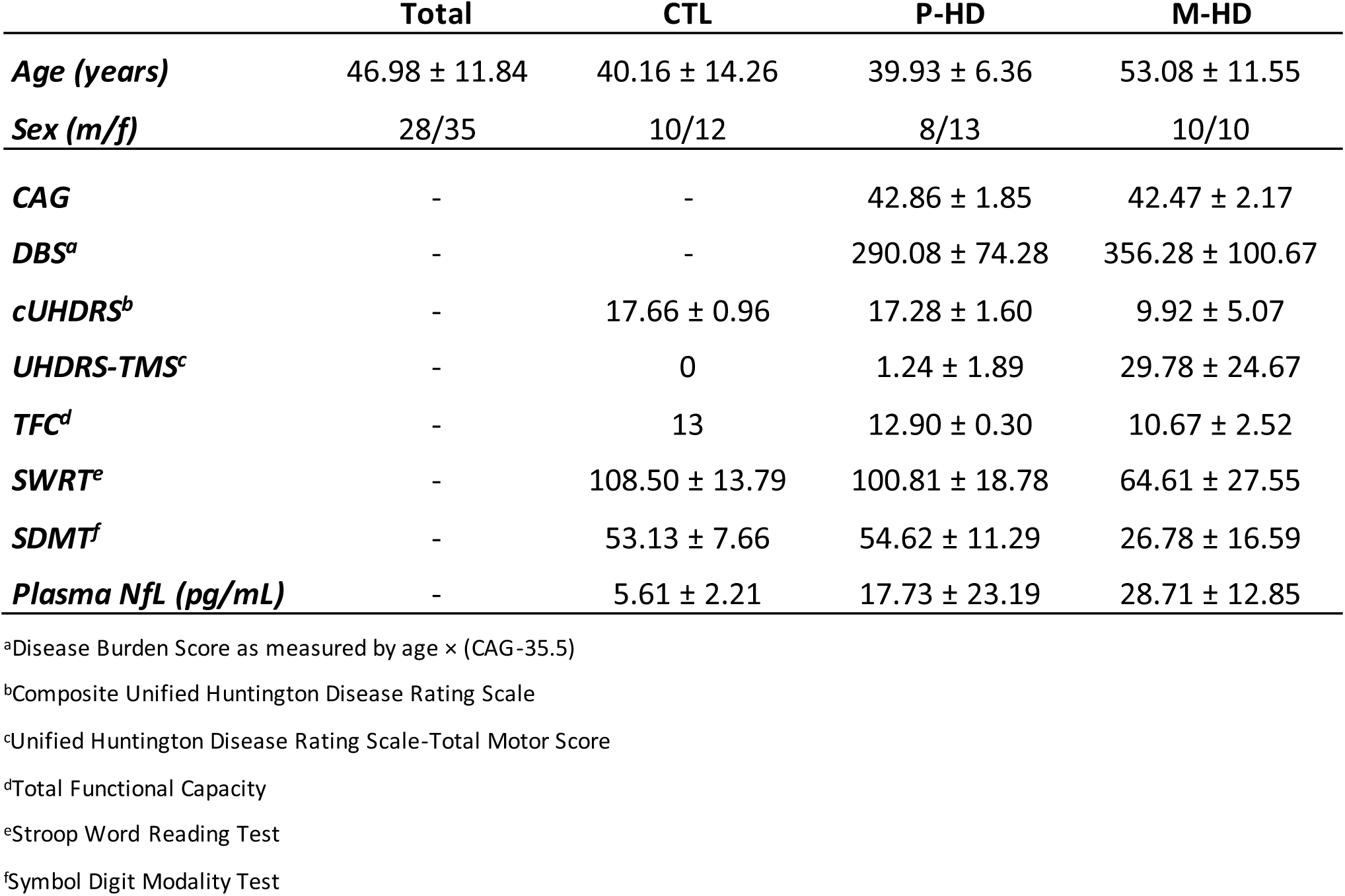
Clinic and sociodemographic characteristics of samples.

|  | Total | CTL | P-HD | M-HD |
| --- | --- | --- | --- | --- |
| **Age (years)** | 46.98 ± 11.84 | 40.16 ± 14.26 | 39.93 ± 6.36 | 53.08 ± 11.55 |
| **Sex (m/f)** | 28/35 | 10/12 | 8/13 | 10/10 |
| **CAG** | - | - | 42.86 ± 1.85 | 42.47 ± 2.17 |
| **DBS<sup>a</sup>** | - | - | 290.08 ± 74.28 | 356.28 ± 100.67 |
| **cUHDRS<sup>b</sup>** | - | 17.66 ± 0.96 | 17.28 ± 1.60 | 9.92 ± 5.07 |
| **UHDRS-TMS<sup>c</sup>** | - | 0 | 1.24 ± 1.89 | 29.78 ± 24.67 |
| **TFC<sup>d</sup>** | - | 13 | 12.90 ± 0.30 | 10.67 ± 2.52 |
| **SWRT<sup>e</sup>** | - | 108.50 ± 13.79 | 100.81 ± 18.78 | 64.61 ± 27.55 |
| **SDMT<sup>f</sup>** | - | 53.13 ± 7.66 | 54.62 ± 11.29 | 26.78 ± 16.59 |
| **Plasma NfL (pg/mL)** | - | 5.61 ± 2.21 | 17.73 ± 23.19 | 28.71 ± 12.85 |

<sup>a</sup>Disease Burden Score as measured by age × (CAG-35.5)
<sup>b</sup>Composite Unified Huntington Disease Rating Scale
<sup>c</sup>Unified Huntington Disease Rating Scale-Total Motor Score
<sup>d</sup>Total Functional Capacity
<sup>e</sup>Stroop Word Reading Test
<sup>f</sup>Symbol Digit Modality Test

### 3. Heterogeneous diversity of extracellular sRNAs in CTL and *HTT* mutation carriers

To uncover the sRNA diversity in the two selected plasma fractions between *HTT* mutation carriers and control samples, we characterized the sRNA profiles by deep sequencing and applied diverse supervised and unsupervised statistical methods to capture differently expressed sRNAs (Fig. 2A). As observed in the proof of concept (Fig. 1F, Supplementary data Fig. 2A), the majority of reads mapped onto rRNA (51%) followed by gene fragments (34%), and a comparable total number of different sRNA clusters were identified in the three groups (Fig. 2I, 2J, Supplementary data Fig. 5A, 5B). However, we observed heterogeneous proportions of some sRNAs biotypes in the two plasma fractions, including miRNAs and tRFs (Fig. 2I, 2J), in line with the initial plasma sub-fractionation analysis (Fig. 1).

It is worth mentioning that sequencing analyses revealed low levels of miRNAs which accounted for less than 5% of the EVs-sRNA content in the three groups of samples (Fig. 2I) as previously shown^31^. The analysis of the NonEV fractions suggest an increased proportion of miRNAs in mutation carriers (Fig. 2J); however, we failed in validating significant changes in qRT-PCR assays (data not shown). This may be related to the significant increase of the inter-sample variability in sRNAs levels in NonEV compared with EV fractions (Fig. 2I, J; Supplementary data Fig. 5C, D), assessed through the comparison of the coefficient of variation (CV) between the two plasma fractions (CV in EVs fraction=0.09; CV in NonEVs fraction=1.1; p-val=0.0027 based on Levene‘s test for homogeneity of variances). Another obvious sign of increased heterogeneity in the NonEV fraction within the three groups of patients is detected in the reads length profiles obtained in the analysis of sequencing data (Supplementary data Fig. 9B).

In contrast, in the EV fractions, probably due to the conferred vesicular protection, the proportions of sRNAs are more homogeneous between individuals (Fig. 2I and Supplementary data Fig. 5C). Thus, we opted to proceed with the downstream analyses only in the vesicular compartment.

### 4. Specific exRNAs-EVs correlate with cognitive and motor performance in HTT mutation carriers

Focusing on EV fraction, through differential expression analysis, our results highlighted a significant downregulation of most of the deregulated sRNAs in mutation carriers compared to CTL samples, with many changes commonly occurring at premanifest stages (Fig. 3A, 3B, Supplementary Table 2, Supplementary Table 3). Most of these deregulated sRNAs were annotated as miRNAs, tRFs, and gene fragments (Fig. 3C). We applied PLS-DA model to highlight specific expression patterns of sRNAs discriminating groups (Supplementary Table 4, Supplementary Table 5). miRNAs, tRFs, and gene fragments that contributed more to the PLS-DA discrimination, (showing a variable importance in projection –VIP-value > 0.8) were selected for unsupervised principal component analysis (PCA). The first two components of the PCA showed that P-HD and M-HD clearly separated from healthy controls (Fig. 3D), while separation was not so obvious between P-HD and M-HD, which could be achieved when using sRNAs identified with the ExceRpt tool (Supplementary data Fig. 6D).

**Figure 3:**
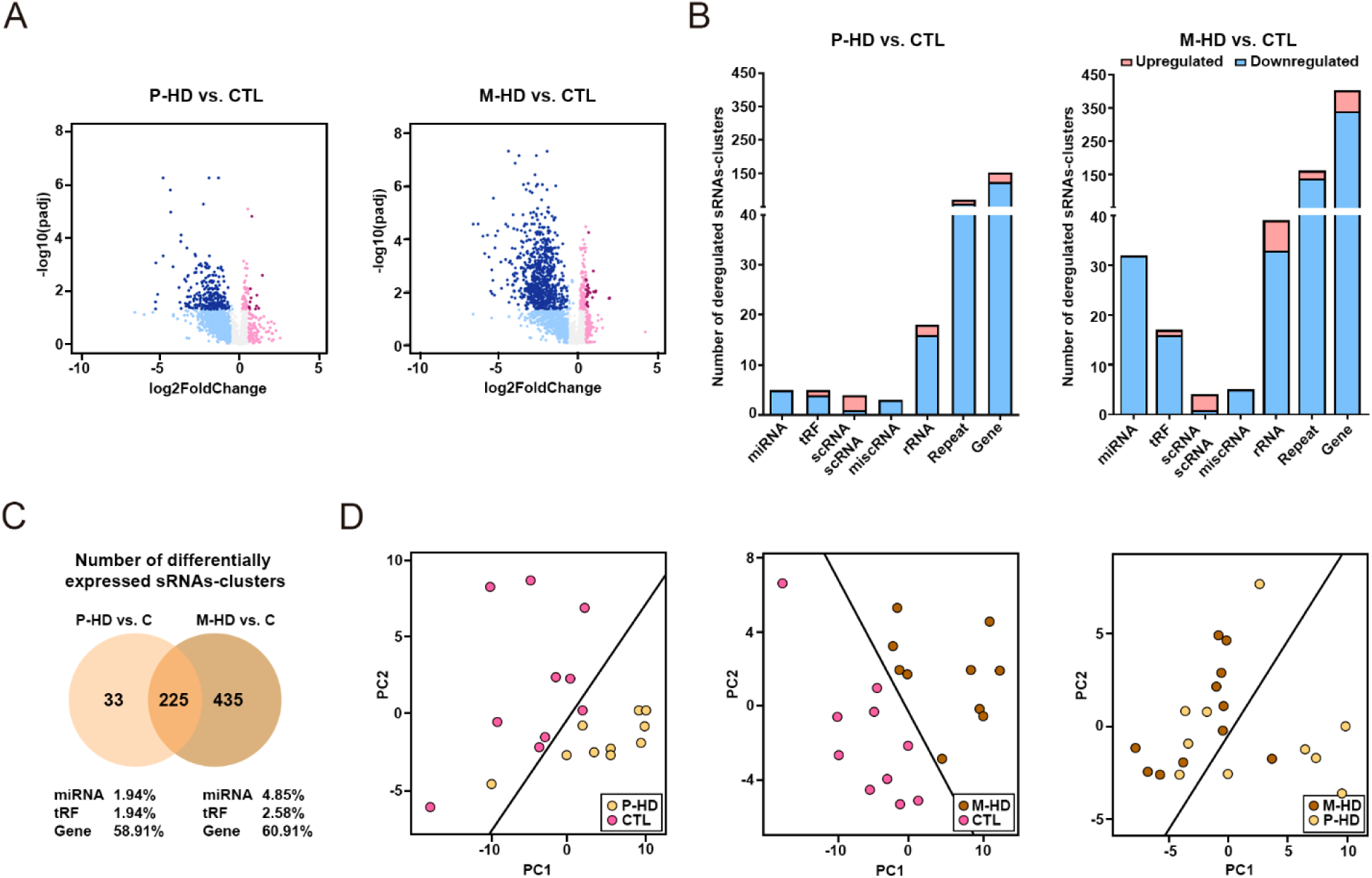
sRNAs from HD patients are deregulated in EVs fractions in comparison to CTL samples, using SeqCluster tool. (A) Volcano plots showing differentially expressed sRNAs-clusters in P-HD-EVs versus CTL-EVs fractions and M-HD-EVs versus CTL-EVs fractions. Orange dots represent significantly deregulated genes (|log2FoldChange|>0.58, adjusted P<0.05, n= 10 per group). (B) Total number of sRNAs-clusters differentially expressed between P-HD-EVs versus CTL-EVs fractions and M-HD-EVs versus CTL-EVs fractions. (C) Venn Diagram of differentially expressed sRNAs-clusters between P-HD vs CTL and M-HD vs CTL, showing the number of overlapped dysregulated sRNAs-clusters between both comparisons. (D) PCA plots constructed with top DE sRNAs and top sRNA that contribute to discriminate between disease conditions and healthy individuals based on PLS-DA (VIP > 0.8). The solid line is the linear discriminant function that best separates disease conditions.

To gain insight into the association of differentially expressed (DE) sRNAs (miRNAs, tRFs, and gene fragments) and the disease, we explored the correlation between their expression levels and patients’ clinical features considered during the HD course. To define potentially meaningful sets of sRNAs, we applied simple linear regression analysis between each of the components of the cUHDRS (SWRT, SDMT, UHDRS-TMS, and TFC) and individual sRNA-clusters DE in P-HD and/or M-HD versus CTL samples (adj. P < 0.05) and reaching a PLS-DA based VIP > 0.8. This allowed the identification of three smaller sets of sRNAs significantly associated with cUHDRS (p-val < 0.05): 12 DE sRNAs in P-HD, 17 DE sRNAs in P-HD and in M-HD, and 48 sRNAs DE in M-HD (Fig. 4B, Supplementary Table 6). To evaluate the multivariate association between these sets of sRNAs with any of the cUHDRS components, we performed an age– and sex-adjusted PLSR analysis (Table 2). Noticeably, we observed that the expression pattern of the sRNAs specifically DE in P-HD was related to the performance of the SWRT test in *HTT* mutation carriers (P-HD and M-HD) (Fig. 4C, Table 2; coefficient of determination (R^2^) = 0.97; p-val <0.0001). Moreover, when we restricted the analysis to the P-HD group, the association of this set of sRNAs and SWRT was still maintained (Fig. 4D, Table 2; R^2^ = 0.93; p-val = 0.04) while no correlation was observed in the M-HD group (Fig. 4E, Table 2, R^2^ = 0.29; p-val = 0.74), suggesting that the joined expression pattern of a specific set of sRNAs can sense pre-clinical changes. To assess to what extent the correlation observed between SWRT score and the set of top DE sRNAs in P-HD is mainly due to the disease and not a consequence of population diversity, we repeated the analysis including the CTL samples, and observed that the association lost significance (R^2^ = 0.09; p-val = 0.82). The expression levels of this specific set of P-HD DE sRNAs did not show a significant correlation with UHDRS-TMS or TFC in M-HD patients, stressing the specific relevance of these species to highlight premanifest cognitive changes (Table 2).

**Figure 4:**
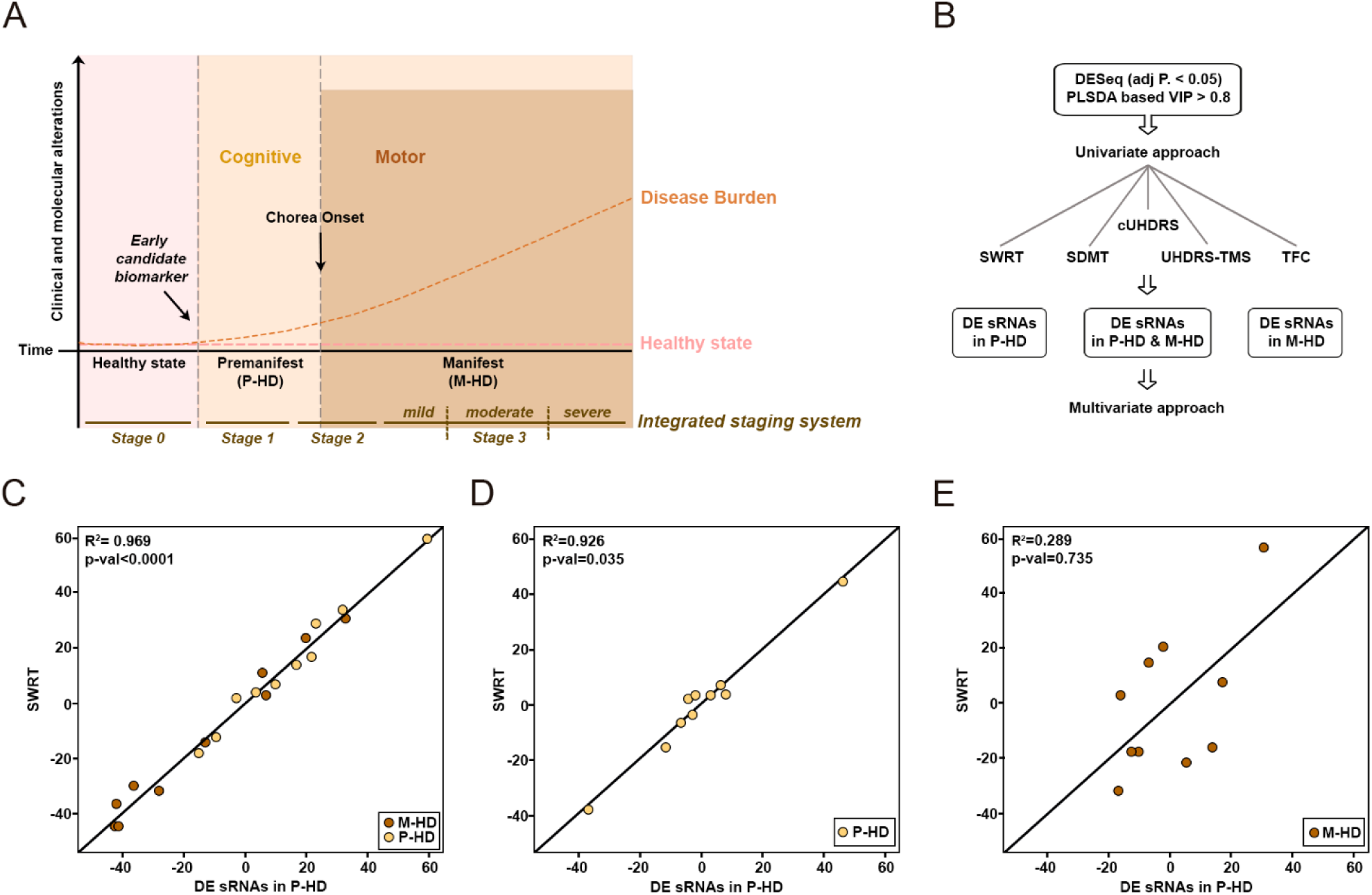
sRNAs deregulated in P-HD correlate with cognitive symptomatology. (A) Framework of Huntingtońs Disease stages. The ideal candidate molecular biomarker would be measurable prior to the appearance of P-HD and M-HD symptoms and would correlate with the disease course. The current patients’ classification system together with the Integrated staging system ^57^ are depicted. (B) Workflow followed to select relevant sRNAs for correlation analyses. PLSR analysis depicting the relation between the set of sRNAs specifically DE in P-HD patients and the performance of the SWRT test (C) in HTT mutation carriers (P-HD and M-HD; R^2^=0.97; p-val<0.0001); (D) only in the P-HD group (R^2^=0.93; p-val=0.035); and (E) only in the M-HD group (R^2^=0.289; p-val=0.735).

**Table 2:** Correlation of EV DE sRNAs and clinical measures in HTT mutation-carriers.

|  | Groups included | SWRT | SDMT | UHDRS-TMS | TFC | cUHDRS |
| --- | --- | --- | --- | --- | --- | --- |
| **sRNAs DE in P-HD (12)** | P-HD, M-HD (n=20) | **0.97 (p-val< 0.0001)** | 0.16 (p-val: 0.62) | - | - | 0.19 (p-val: 0.79) |
|  | P-HD (n=10) | **0.93 (p-val: 0.04)** | 0.38 (p-val: 0.72) | - | - | 0.65 (p-val: 0.24) |
|  | M-HD (n=10) | 0.29 (p-val: 0.74) | 0.29 (p-val: 0.58) | 0.297 (p-val: 0.79) | 0.37 (p-val: 0.79) | 0.28 (p-val: 0.88) |
| **sRNAs DE in P-HD & M-HD (17)** | P-HD, M-HD (n=20) | 0.31 (p-val: 0.51) | 0.34 (p-val: 0.25) | - | - | 0.41 (p-val: 0.13) |
|  | P-HD (n=10) | 0.62 (p-val: 0.41) | 0.59 (p-val: 0.30) | - | - | 0.42 (p-val: 0.79) |
|  | M-HD (n=10) | 0.59 (p-val: 0.18) | 0.50 (p-val: 0.18) | 0.44 (p-val: 0.39) | 0.53 (p-val: 0.23) | 0.56 (p-val: 0.17) |
| **sRNAs DE in M-HD (49)** | P-HD, M-HD (n=20) | 0.24 (p-val: 0.97) | 0.15 (p-val: 0.99) | - | - | 0.17 (p-val: 1) |
|  | P-HD (n=10) | 0.39 (p-val: 0.99) | 0.06 (p-val: 1) | - | - | 0.18 (p-val: 1) |
|  | M-HD (n=10) | **0.85 (p-val: 0.03)** | 0.69 (p-val: 0.07) | **0.79(p-val: 0.004)** | 0.46 (p-val: 0.93) | **0.89 (p-val < 0.0001)** |

Partial least squares regression age-adjusted was conducted. cUHDRS: Composite Unified Huntington Disease Rating Scale; UHDRS\_TMS: Unified Huntington Disease Rating Scale-Total Motor Score; TFC: Total Functional Capacity; SWRT: Stroop Word Reading Test; SDMT: Symbol Digit Modality Test.

Parallel regression analyses with the set of sRNAs specifically DE in M-HD patients showed significant correlations with SWRT, SDMT, UHDRS-TMS, and cUHDRS parameters (R^2^ = 0.85, p-val = 0.03; R^2^ = 0.69, p-val = 0.07; R^2^ = 0.79, p-val = 0.004; and R^2^ = 89, p-val < 0.0001, respectively) when considering only the manifest group (Table 2). Of notice, the association was lost when the P-HD and CTL groups (UHDRS-TMS score > 5) were included in the analysis. Overall, these data suggest that the combined levels of specific sets of sRNAs can sense pre-clinical changes and/or symptoms.

### 5. Sensitivity and specificity of candidate biomarkers for premanifest HD

As potential new biomarkers in HD and aiming to validate the sRNAs-sequencing results, eight sRNAs candidates (DE sRNAs showing a p-adj < 0.05 in P-HD vs CTL and in M-HD vs CTL, with a base mean sequence reads > 200 and PLSDA based VIP > 0.8) were selected for qRT-PCR determinations. We validated their relative expression in 20 samples per group, including the same set of samples used in sRNA sequencing (n=10 samples/group) and in an independent biological set of samples (n=10 samples/group). Due to the absence of universal endogenous sRNA normalizers, hsa-miR-100-5p was selected as the reference gene since it was a highly stable sRNA with low variability among all the sequenced samples (Supplementary data Fig. 7).

According to qRT-PCR results, miR-451a, miR-21-5p, miR-26a-5p, let-7a-5p, miR-27a-3p, tRF-Glu-CTC, and tRF-Gly-GCC expression were significantly downregulated in P-HD and M-HD in comparison to CTL samples, consistent with the sequencing findings. tRF-Lys-TTT expression showed an analogous downregulation between M-HD versus CTL samples, considering an unadjusted p-value of 0.015 (Fig. 5A).

**Figure 5:**
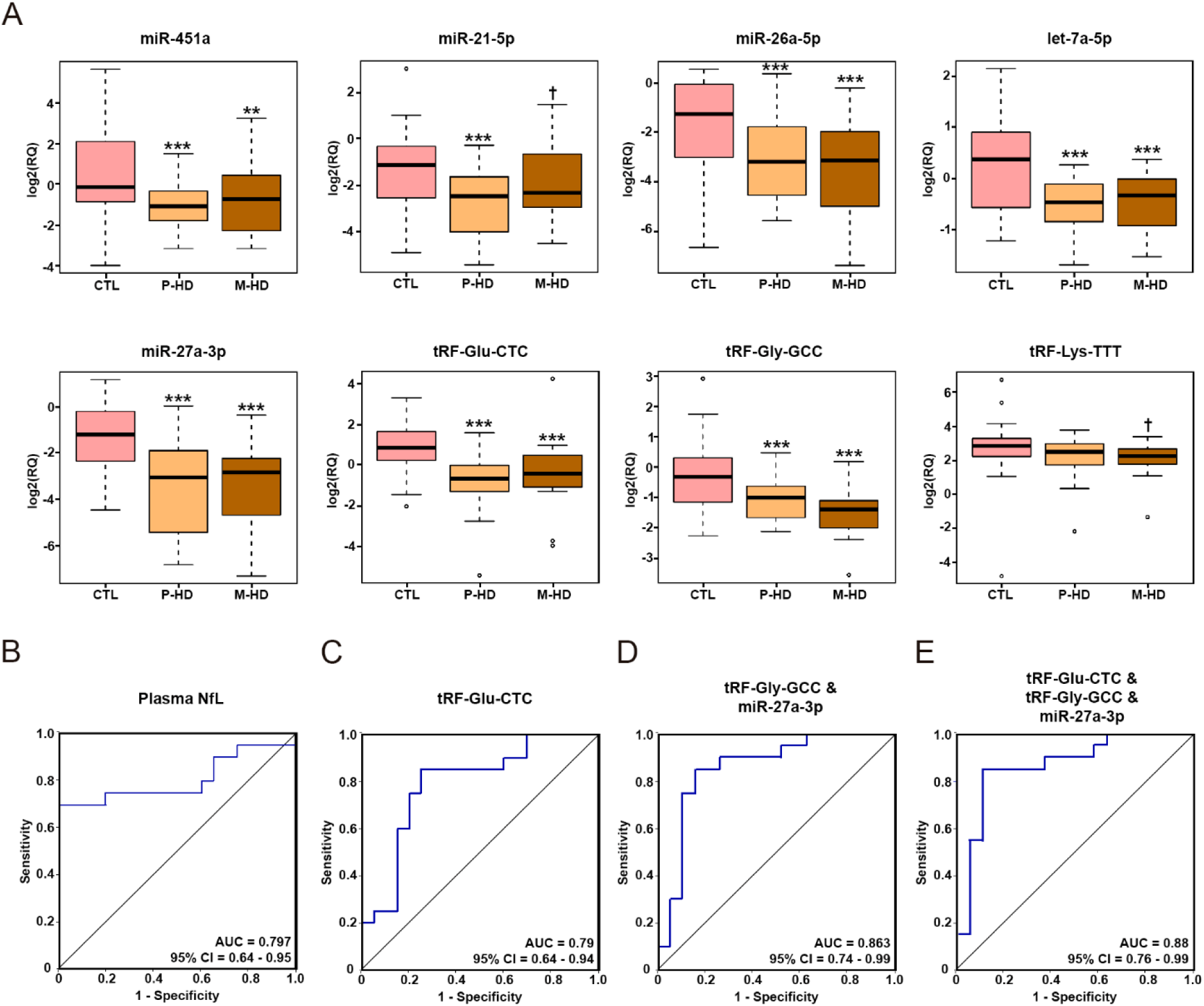
Validation and diagnostic potential analysis of selected sRNAs at premanifest stages. (A) Boxplots representing the relative expression of sRNAs validated with qRT-PCR in technical and biological groups of samples. Significant differences between P-HD and M-HD versus CTL groups are presented with *** (adj P <0.001), ** (adj P <0.01) and nominally significant differences are presented with **ⴕ** (p-val < 0.05). (B) ROC curves analysis of the sensitivity and specificity of plasma NfL between P-HD and CTL (p-val=0.001). (C) Representative ROC curve analysis of the sensitivity and specificity of an individual validated sRNA: tRF-Glu-CTC between P-HD and CTL (p-val=0.002). (D) ROC curves analysis of the sensitivity and specificity of a novel 2-sRNAs-biosignature: ensemble of tRF-Gly-GCC and miR-27a-3p between P-HD and CTL (p-val=0.0001). (E) ROC curves analysis of the sensitivity and specificity of a novel 3-sRNAs-biosignature: the ensemble of tRF-Glu-CTC, tRF-Gly-GCC and miR-27a-3p between P-HD and CTL (p-val=0.0001). n=20 per group.

To evaluate the sensitivity of the qRT-PCR validated sRNAs as biomarkers for early HD stages, we analyzed their biomarker prediction outcome. At the same time, we analyzed the diagnostic potential of NfL in the same group, as it is considered the most sensitive blood biomarker in HD (Fig. 5B). sRNAs individually showed a similar diagnostic potential compared to NfL (Fig. 5C, Supplementary data Fig. 8A). Of note, our data indicated that the combination of two, three, or four validated sRNAs showed similar specificity and sensitivity discriminating between P-HD patients and healthy individuals (Fig. 5D, 5E, Supplementary data Fig. 8B). For instance, the performance of an ensemble of 2-sRNAs-biosignature including tRF-Gly-GCC and miR-27a-3p presented an AUC = 0.863 (CI = 0.74-0.99, Fig. 5D), and the ensemble of 3-sRNAs-biosignature being tRF-Glu-CTC, tRF-Gly-GCC and miR-27a-3p, reached an AUC = 0.88 (CI = 0.76 – 0.99, Fig. 5E).

Likewise, using these sets of patients and considering the use of plasma NfL in combination with our validated sRNAs, we also estimated a high diagnosis potential. Specifically, the combined biosignature of tRF-Gly-GCC and miR-27a-3p, together with plasma NfL showed an AUC=0.908 (CI = 0.82 – 0.99) (Supplementary data Fig. 8C).

### 6. miR-21-5p as a progression biomarker at premanifest HD stage

Finally, to explore the potential of these sRNAs as progression biomarkers, we analyzed their expression in nine P-HD patients for whom we had paired plasma samples from a 1.5-year follow-up visit (Supplementary Table 1). Specifically, miR-21-5p expression was significantly decreased over time, indicating that it could act as progression biomarker at premanifest stages of the disease (Fig. 6).

**Figure 6:**
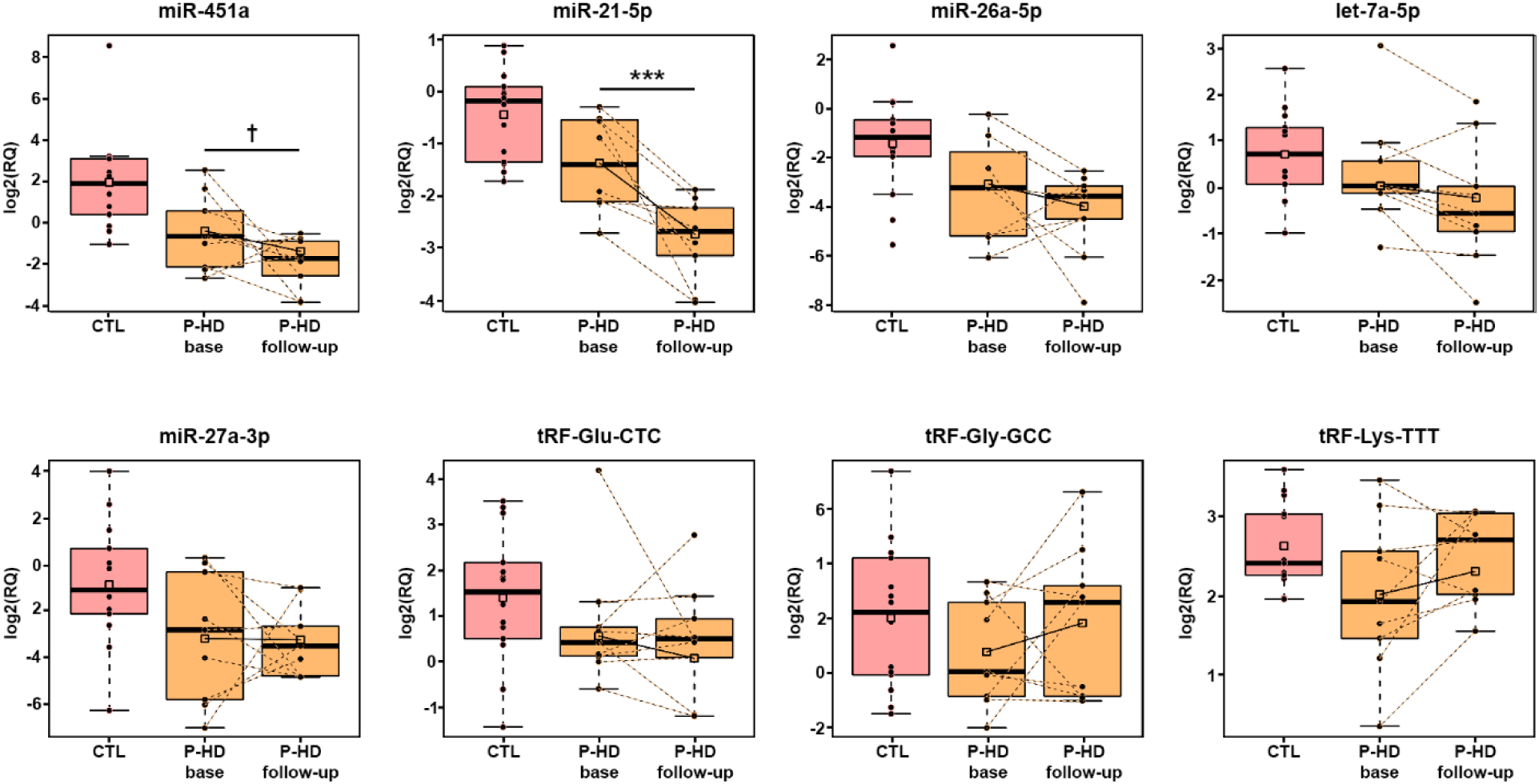
Longitudinal changes validation of selected sRNAs at premanifest stages over a 1.5-year follow-up. Boxplots representing the relative expression of sRNAs validated by qRT-PCR in P-HD paired samples from baseline and 1.5-year follow-up visits. Squares represent mean values. Significant differences between longitudinal samples are presented with *** (adj P <0.001), ** (adj P <0.01), * (adj P <0.05), and nominally significant differences are presented with **ⴕ** (p-val < 0.05).

We further evaluated the correlation between individual sRNAs and the patientś clinical features (Table 3). We observed that the expression of miR-21-5p and miR-26a-5p significantly correlated with SDMT (rho: 0.57, p-val: 0.01) and cUHDRS (rho: 0.65, p-val: 0.002) when only P-HD individuals were considered, supporting these miRNAs as biomarkers sensing preclinical changes. Furthermore, these results suggest that miR-21-5p could be a progression biomarker to monitor longitudinal clinical changes occurring during the premanifest stage. Parallel correlation analyses with NfL and the clinical measures were performed, but significant association was only found when considering the ensemble of P-HD and M-HD groups (Table 3), as previously reported^32,33^.

**Table 3:**
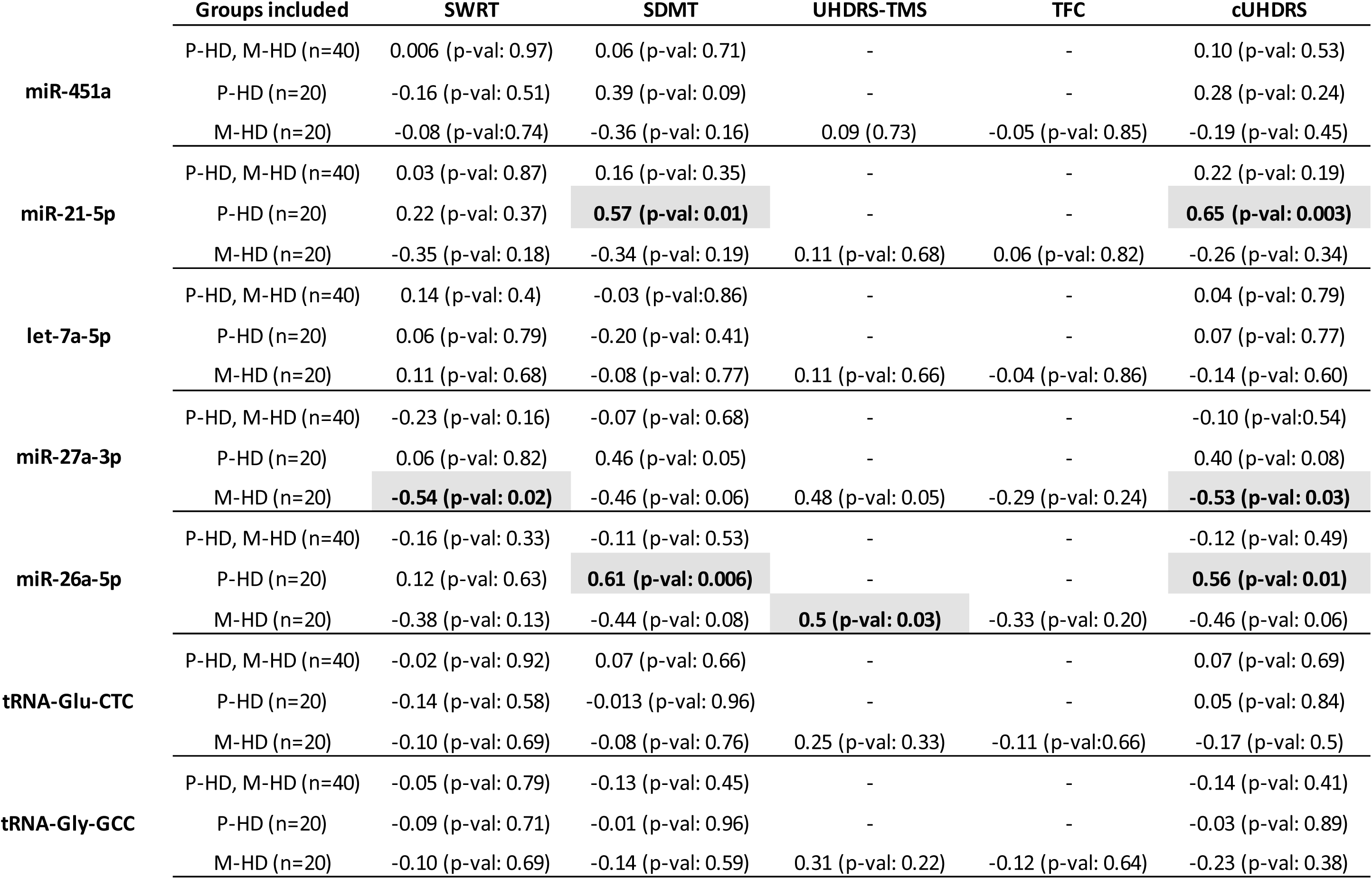
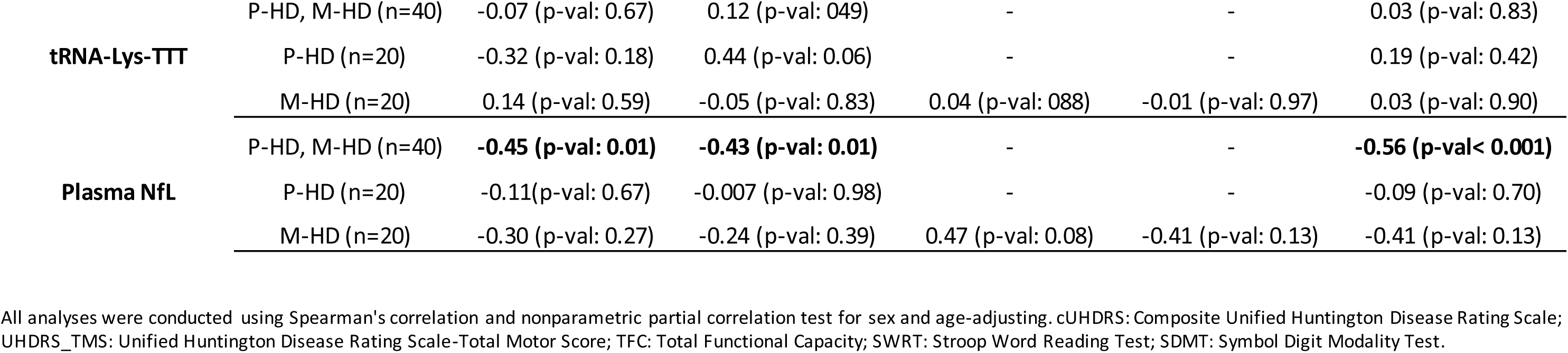
Correlation of EV-sRNAs biosignature as measured by qRT-PCR and plasma NfL levels with clinical measures in HTT mutation-carriers.

## Discussion

In HD, the identification of early molecular alterations in premanifest individuals, occurring before the motor-based clinical diagnosis is of outmost importance, as it provides an opportunity to monitor disease progression and achieve clinical management of the patients in an individual manner. Biofluid sRNAs are a promising source of early biomarkers since their expression levels are highly sensitive to physiological and pathological changes. In HD this is reinforced by the fact that sRNAs are profoundly altered in the brain and these perturbations have been directly linked to HD pathogenesis^17,34,35^. Thus, in addition to clinical measures, sRNA profiling offers opportunities for the direct quantification of pathobiological processes at the molecular level. In the present study, we have deeply analyzed sRNA profiles in plasma sub-fractions and uncovered specific EVs-sRNAs as potential markers to monitor premanifest changes.

A major challenge in reliable and reproducible sRNA biomarker discovery is the identification of an informative biofluid compartment. Here we have identified differentiated sRNA profiles in EVs (cEVs) and extravesicular, protein enriched (NonEVs-High) plasma subfractions, confirming the idea that diverse plasma carriers offer scenarios for differential distribution of exRNAs^36^. However, it became apparent that cEVs were a more suitable source to analyze sRNAs as biomarkers, showing greater stability compared with NonEVs-High fraction. As widely discussed elsewhere^37,38^, EV-sRNAs are highly resistant to degradation, which may contribute to more consistent and uniform exRNA profiles.

In HD, several studies have been focused on the exploration of brain and CSF miRNAs expression^39–41^. However, little information is available regarding dynamic changes of sRNAs in plasma, and none has studied plasma-EVs as a source of biomarkers. Furthermore, while exRNA biomarker research is mainly focused on miRNAs we showed that miRNAs represent a minor proportion of the EVs transcriptome, in line with previous reports^31,42–45^, and other species including tRFs and YRNAs are highly represented.

Here, we have shown that most of the plasma EVs-derived sRNAs are downregulated in mutation carriers compared with CTL, with more significant changes at manifest stages. Previously reported upregulated miRNAs in plasma of HD patients^46,47^ including miR-10b-5p, miR-486-5p, miR-30d-5p, and miR-222-3p were not validated in the present study. In addition, the here observed downregulated miR-27a-3p and miR-21-5p were significantly overexpressed in analyses of total plasma as reported by Díez-Planelles and colleagues^46^. Overall, these data suggest that the specific EVs miRNA content may offer complementary information regarding HD status.

The massive downregulation of sRNAs observed at premanifest and manifest stages in HD –EVs might reflect an altered vesicular sRNA-packaging mechanism. Currently, the specific pathways involved in the RNA packaging in EVs remain unknown^48^. However, studies are increasingly revealing that active sorting and loading of coding and non-coding RNAs in EVs may be controlled by RNA-binding proteins. These include the hnRNP family^49,50^ and/or via ESCRT dependent or independent pathways^51^. Whether alterations of these proteins could lead to perturbations in the RNA content of HD EVs remains to be answered.

Our study shows that beyond miRNAs, tRF-Gly-GCC and tRF-Glu-CTC are early altered in mutation carriers. These species have been recently revealed as highly abundant and stable in circulation^52,53^, pointing them as promising pre-manifest biomarkers in HD. In addition, in HD, strong deregulation of these species, including the EV-validated tRF-Gly-GCC and tRF-Glu-CTC, has been detected in the putamen of patients^17^, and functional analysis suggests a role of diverse tRFs in neuronal dysfunction^17^. Future studies should determine whether the dynamic expression of circulating tRFs reflects brain alterations.

HD clinical alterations emerge gradually during a premanifest phase, and the precise shift from premanifest to manifest HD may be difficult to anticipate (Fig. 4A). Subtle cognitive changes can be detected in premanifest population up to 15 years before diagnosis in specific measures addressing processing speed and attention^54^. This is the case of the SWRT whose linear worsening has been extensively reported to occur from the premanifest stage of the disease. Based on sequencing data analysis, we have observed that the expression of a specific set of sRNAs DE in P-HD (and not in M-HD) was strongly associated with SWRT score using the P-HD group (either individually or with M-HD), but not with the UHDRS-TMS in the M-HD group. Complementing this finding, a set of sRNAs specifically DE in M-HD significantly correlated with UHDRS-TMS and cognitive performance (SWRT and SDMT) when evaluating the M-HD group, but not with the P-HD samples, either alone or together with M-HD. These data indicate that specific sets of sRNAs may distinguish premanifest alterations and manifest symptomatology, pointing the as highly sensitive biomarkers.

The translational utility of these findings is limited by the sensitivity of the current techniques (qRT-PCR) to detect low-abundant sRNAs and the type of candidate sRNAs. Several sRNAs within the specific sets that significantly correlate with clinical signs correspond to gene fragments. Many of the altered gene fragments may provide an indirect measure of the levels of changing genes rather than bona fide sRNAs that can be proposed as reliable molecular biomarkers. Therefore, subsequent follow-up validation in additional samples focused on genuine sRNAs such as miRNAs and tRFs consistently detected and significantly DE according to sequencing data analysis. The successful validation of 7 out of 8 distinct sRNAs candidates by qRT-PCR in independent samples confirmed the robustness of our sRNA-seq analysis pipeline. Interestingly, the follow-up validation experiment in a group of P-HD patients revealed the potential of miR-21-5p as pre-clinical progression biomarker. Of notice, when using P-HD samples individually, miR-21-5p and miR-26a-5p were significantly correlated with SDMT, which gradually indexes disease progression at these pre-motor stages^55^, and with cUHDRS. On the other hand, we failed to obtain a significant correlation between NfL plasma levels and clinical data in this set of premanifest samples, which was only reached when using P-HD and M-HD groups together, validating what has recently been published^32,33^. NfL is rising as a reference biomarker reflecting neuronal injury in multiple neurodegenerative disorders. However, the present results suggest that NfL presents a limited utility as a tracking measure for premanifest changes, which could be overcome by analyzing specific sRNAs.

Confirming previous studies^32^, our results indicate that Nfl levels could strongly distinguish M-HD from both P-HD and Control groups. Candidate sRNAs and NfL displayed a similar discriminative performance between P-HD or M-HD and CTLs; however, sRNAs could not differentiate between P-HD and M-HD groups. Overall, these data indicate a clear progression of neuronal damage and axonal injury from premanifest to manifest stages (Nfl levels) that was not mirrored by the selected candidate sRNAs, strongly altered at both stages. Nevertheless, the robust alteration of specific sRNAs at premanifest stages suggests that these species may finely sense early perturbations occurring long before obvious neurodegeneration. Supporting this hypothesis, changes in gene expression at the transcriptional level (both mRNA and miRNA) occur in the brain of HD mouse models before the onset of motor and cognitive alterations, and striatal cell loss^35,56^.

Thus, studying sRNAs that reflect changes occurring before the onset of neurodegeneration and clinical symptoms, appears to be a promising approach to better classify mutation carriers at premanifest stages of HD. Specifically, this approach could aid in the stratification of individuals in stage 0 to stage 1 of the Integrated Staging System for the classification of HD (as shown in Fig. 4A), which was developed to address the need for better subclassification criteria in the pre-motor HD stages^57^.

Despite these findings, there are still several limitations in our study. Notably, the clinical implementation of RNA-based biomarkers in EVs demands rigorous analytical validation and standardized procedures for EV isolation, RNA extraction, and detection of specific species. These requirements are mandatory to ensure the reliability and applicability of EV-based biomarkers in clinical settings. In addition, we acknowledge the risk of over-fitting RNAseq for biomarker discovery. While we have conducted in-depth analysis of sRNA profiles in a discovery cohort and validated specific candidates in a larger group of patients, further confirmatory studies in multi-center settings with larger cohorts are necessary. Moreover, the relatively short follow-up time does not allow interpretation of our findings in relation to significant aggravation of clinical symptomatology.

In summary, our results demonstrate that, among the heterogenous sRNA plasma composition, encapsulated EV-sRNAs are early deregulated in HD, and that this deregulation might be associated with early changes occurring at pre-motor phases of the disease. Altogether, these findings present a compelling argument to consider specific sRNAs, including diverse biotypes such as tRFs, as promising HD biomarkers.

## Methods

### Participants

In this study, all participants were recruited from the Movement Disorders Unit at Hospital de la Santa Creu i Sant Pau (Barcelona, Spain). We included forty-one Caucasian gene mutation carriers and twenty-two Caucasian healthy non-mutation carrier control participants (Table 1, Supplementary Table 1). Individuals were confirmed gene mutation carriers (CAG repeat length > 40), and disease – specific indicators including motor, cognitive, and neuropsychiatric status, were recorded.

The disease burden score (DBS) was used as an index of pathological burden due to lifetime exposure to mutant huntingtin and calculated as age (in years) x (CAG – 35.5). Gene-mutation carriers were grouped into premanifest (P-HD, n=21) and manifest (M-HD, n=20) stages based on the Unified Huntington’s Disease Rating Scale total motor score (UHDRS-TMS) and the Total Functional Capacity (TFC). Participants were classified as M-HD patients if UHDRS-TMS > 5 and a diagnostic confidence level=4, indicating that motor abnormalities were unequivocally caused by HD with ≥99 % of confidence. Participants with a diagnostic confidence level (DCL) < 4 and a TFC = 13 were considered P-HD.

The following cognitive indicators, known to be sensitive to HD progression, were also administered: The Symbol Digit Modality Test (SDMT) and the Stroop-word reading test (SWRT)^58,59^. The Composite UHDRS (cUHDRS) score was also calculated^60^.

All participants provided written informed consent previously approved by the appropriate ethics committee. The study was therefore performed in accordance with the ethical standards laid down in the 1964 Declaration of Helsinki and its later amendments.

### Sample collection and processing

Blood samples were collected in tubes with 0.109 M 3.2% trisodium citrate (Vacutainer, Becton Dickinson) via antecubital vein puncture in the morning, and processed within 30 min after withdrawal. Tubes were centrifuged at 500 g for 10 min at room temperature (RT) and the supernatant was pipetted off from cell debris. The supernatant was centrifuged again at 2,500 g for 15 min at RT to eliminate platelets and one more time at 13,000 g for 15 min at RT to eliminate bigger particles and smaller aggregates. Platelet-free plasma was collected and stored directly at –80°C.

### Plasma NfL

Neurofilament light (NfL) was measured with the Simoa Human NF-light Advantage kit (Catalogue # 103400) using the Single Molecule Array (Simoa) technology (Simoa; Quanterix, Lexington, MA, United States) in the SR-X Biomarker detection system by following the manufacturer’s instructions. Briefly, the calibration curve, control samples, inter-assay control (pooled plasma with known concentration), and unknown samples in duplicate were loaded onto a 96-well plate. Samples were incubated with antibody-coated paramagnetic beads and a biotinylated antibody detector simultaneously. After a wash, streptavidin-conjugated β-galactosidase (SBG) reagent was added to bind the biotinylated antibodies, leading to the SBG enzyme labeling of the capture of NfL. Once in the SR-X platform, the beads were resuspended in resorufin β-D-galactopyranoside (RGP) reagent, transferred to the Simoa disk and sealed. The NfL proteins captured by the antibody-coated paramagnetic beads and labeled with the SBG reagent hydrolyze the RGP substrate to produce a fluorescent signal. The fluorescent signal values generated from the calibration curve of known concentrations were fit using a 4-parameter logistic curve that was used to calculate the unknown and control sample concentrations. All the samples were measured in duplicate and had an intra – assay coefficient of variation (CV) <20% with a mean of 7.54%. Inter –assay CV% was 9.34%.

### Plasma fractionation and EVs isolation

For plasma fractionation and EVs isolation, we used size-exclusion chromatography (SEC) followed by ultrafiltration (UF) as previously described^25,26^. Briefly, Sepharose CL-2B (Sigma Aldrich, MO, USA) was stacked in Puriflash Dry-Load Columns (Interchim) of 20 mL. On top of each column, two mL of plasma were loaded, and fractions of 500 μL were collected manually in individual tubes (a total of 35 tubes-fractions) using filtered 1X PBS as elution buffer. Protein content for each fraction was measured by Micro BCA Protein Assay Kit (Thermo Fisher Scientific) and Bio-Rad Protein Assay Dye Reagent (BioRad). SEC-fractions were further characterized using classical EVs-associated markers CD9, CD63 and CD81 by bead-based flow cytometry assay as previously reported^24^ in a BD LSRFortessa (BD Biosciences). Median fluorescence intensity (MFI) values were obtained using Diva Software, and the four fractions with the highest MFI compared to the isotype control (fold change MFI) were considered EVs-enriched fractions and were pooled. Purified pooled EVs were further concentrated by ultrafiltration using Amicon Ultra 100 KDa Centrifugal Filters (Merck Millipore) according to the manufacturer’s instructions, obtaining the concentrated EVs (cEV fraction) and Filtrate fractions (Fig. 1A). Pooled fractions were kept at –80°C until later use.

For the determination of sRNA diversity in the different plasma subfractions, four SEC fractions depleted of EVs with low protein concentration (NonEVs-Low fraction) and four SEC fractions depleted of EVs with high protein content (NonEVs-High fraction) were also pooled (Fig. 1B).

For the analysis of expression patterns of plasma sRNAs in HD, eight EV-depleted SEC fractions (identified as NonEVs fraction) were pooled to investigate the extravesicular fraction in the patients’ and healthy controls’ groups (Fig. 2B).

### Nanoparticle tracking analysis

The particle size distribution and concentration measurements in cEVs fractions were evaluated by nanoparticle tracking analysis (NTA) using a NanoSight NS300 instrument (Malvern Panalyitical) and following the manufacturer’s instructions. Each sample was diluted in sterile and filtered 1X PBS to the working range of the system (106–109 particles/mL). Data acquisition and processing were performed using the Nanosight NS300 software (version 3.4) recording three 30-second videos with 25 frames per second. Camera type was sCMOS with Laser Type Blue488 and Camera Level 12. Video analysis settings were set to a detection threshold of 5.

### Western Blotting

Assessment of specific EV markers and Non-EV markers was performed by Western blotting. Protein concentration was measured by Micro BCA Protein Assay Kit (Thermo Fisher Scientific) in the cEVs and Filtrate fractions, samples with low input of protein. Protein concentration in NonEVs-Low fractions, NonEVs-High fractions, and cell lysate samples was determined using Bio-Rad Protein Assay Dye Reagent (BioRad). Equal protein amount of each sample (20 μg) was mixed with reducing 5X Pierce Lane Marker Reducing Sample Buffer (Thermo Fisher Scientific) and boiled for 5 min at 95°C. Due to their not-existing protein content, for Filtrate fractions, the same volume as the one corresponding to 20 ug of the cEVs fractions (from original sample), was loaded. Samples were resolved in NuPAGE 4–12% Bis-Tris polyacrylamide gels (Thermo Fisher Scientific) using MOPS SDS as running buffer (Thermo Fisher Scientific). Gels were transferred to nitrocellulose membranes with the iBlot2 system (Thermo Fisher Scientific). In order to visualize the depletion of Albumin and IgGs in cEVs fractions, Ponceau S staining (Sigma Aldrich) was applied. Membranes were blocked with 5% non-fat dry milk (sc-2325, Santa Cruz Biotech) diluted in 1X Tris-buffered saline with 0.1% Tween 20 (TBS-T) for 1 h at RT. Blots were incubated overnight at 4 °C with the following primary antibodies: rabbit anti-CD9 (13174, Cell Signalling Technology), rabbit anti-Alix (12422-1-AP, Proteintech), mouse anti-Flotillin1 (610820, BD Biosciences), rabbit anti-TSG101 (ab30871, Abcam), rabbit anti-Syntenin (ab133267, Abcam), rabbit anti-Calnexin (ab22595, Abcam) or mouse anti-OxPhos Complex IV subunit IV (anti-COX IV; A-21347, Thermo Fisher Scientific) diluted 1:1000 in TBS-T with 5% Bovine Serum Albumin (BSA, Sigma). Membranes were then washed three times in TBS-T and incubated for 1 h at RT with goat anti-rabbit or goat anti-mouse HRP conjugated secondary antibodies (31460 and 31430, respectively, Thermo Fisher Scientific) diluted 1:10000 in TBS-T. Blots were washed again three times in TBS-T and detected with SuperSignal West Pico Plus or SuperSignal West Femto Maximum Sensitivity chemiluminescent substrates (Thermo Fisher Scientific). Images were acquired in the imaging system Chemidoc (BioRad).

### Cryogenic transmission electron microscopy

Plasma subfractions were imaged by cryogenic transmission electron microscopy (cryo-EM). Vitrified specimens were prepared by placing 3 μL of a sample on a Quantifoil 1.2/1.3 TEM grid, blotted to a thin film, and plunged into liquid ethane-nitrogen in the Leica EM-CPC cryo-work station. The grids were transferred to a 626 Gatan cryo-holder and maintained at −179 °C. The grids were analyzed with a Jeol JEM 2011 transmission electron microscope operating at an accelerating voltage of 200 kV. Images were recorded on a Gatan Ultrascan 2000 cooled charge-coupled device (CCD) camera with the Digital Micrograph software package (Gatan). Size distribution of EVs was quantified by measuring the diameter of around 200 vesicles in each sample using ImageJ.

### RNA extraction and small RNA sequencing

sRNA was isolated using the miRNeasy Serum/Plasma Advanced Kit (Qiagen) according to the manufactureŕs instructions. All samples were adjusted to an input final volume of 200 µL. RNA samples were eluted in 20 µL of RNAse-free water. Prior to sequencing, RNA was precipitated using glycogen (20 mg/ mL, Roche), sodium acetate 3M pH 4.8 (Sigma-Aldrich), and 100% ethanol. After 24 h at –20°C, the mixtures were centrifuged at 14,000 rpm for 30 min at 4°C. Carefully, the supernatant was removed, and the pellet was then washed with 75% ethanol followed by centrifugation at 12,000 rpm for 5 min at 4°C. Finally, the supernatant was discarded, and the RNA pellet was dried at RT and dissolved at 65°C for 3 min in 6 µL of RNAse-free water and stored keeping the cold chain at –80°C.

The RNA samples were analyzed for RNA quality and level of degradation using the Bioanalyzer 2100 (Agilent). sRNA libraries were generated using the NEBNext Small RNA Library Preparation Set for Illumina (New England Biolabs) following the manufacturer’s instructions. Indexed libraries were equimolarly pooled, and the selected band of interest (145-160 bp) was purified. Single-end sequencing of libraries was then carried out on the Illumina HiSeq 2000 with read lengths of 50 nt, generating at least 8 million reads per sample.

### Bioinformatic and biostatistical analysis

The quality of the sequenced fastq files was checked using the FastQC software (http://www.bioinformatics.babraham.ac.uk/projects/fastqc/) and sequence quality scores across all bases of EVs and NonEVs fractions per group were obtained (Supplementary data Fig. 9A). Adapter trimming was performed with Cutadapt^61^ and the trimmed reads were aligned to the human genome (version Ensemble hg19) using STAR aligner (version 2.7.9a)^62^ (Supplementary data Fig. 9B). The obtained sequences were annotated and quantified using SeqCluster ^28^ and ExceRpt (NIH ExRNA Communication Consortium –ERCC-)^29^ bioinformatic tools. During the pre-processing step, ExceRpt filters out highly abundant sRNAs mapping onto ribosomal RNAs (rRNAs) and makes an oversimplification of the highly complex tRNA-derived fragments (tRFs) landscape by only considering the type of isoacceptor. Because non-randomly generated rRNA fragments and tRFs are bioactive molecules^63^ that are emerging as appealing biomarkers^64,65^, we used SeqCluster tool to highlight these biotypes. SeqCluster uses a heuristic iterative algorithm to deal with multi-mapped events. Sequences are consistently and non-redundantly grouped if they map onto the same genomic site defining “clusters” of co-expressed sRNAs. Clusters are then classified according to the type of precursor where sRNAs map, including rRNA-and tRF-clusters. We generated ExceRpt and SeqCluster count matrices with raw reads of each sRNA biotype in every sample. In order to detect outlier samples in our RNAseq analysis, we generated PCA plots with the normalized sRNAs sequence count matrix and we didńt identify any outlier (Supplementary data Fig. 9C).

Prior to further analyses, only sRNAs identified with a minimum of ten counts in at least half of the samples of the tested conditions were considered. Differential expression analyses were performed through negative binomial generalized models implemented in the Bioconductor package DESeq2 ^66^ from R statistical software. Age and sex were considered as confounding factors and Benjamini-Hochberg correction for multiple testing was applied to control the False Discovery Rate.

Partial least squares discriminant analysis (PLS-DA), a multivariate supervised method, based on orthogonal transformations of the initially quantified sRNAs into few uncorrelated components in a way that maximizes the separation of each disease condition, was performed as previously described^67^. Three PLS-DA models were applied for each of the three sRNA biotypes considered (miRNAs, tRFs, and gene fragments). To elude over-parameterized models with rather poor discriminant properties, we obtained in the first stage of the process the most important discriminant clusters from the three PLS-DA models. And, in the second stage, we conducted another PLS-DA analysis including the important clusters associated independently to the three different models found in the first stage. Variable importance for the projection (VIP) criterion was used to identify which sRNAs contribute most to the classification performance, and sRNAs with PLS-DA-based VIP > 0.8 in the second stage were considered for further downstream analysis. Principal Component Analysis (PCA) was applied as a non-supervised method to evaluate the separation between disease conditions using the top contributing sRNAs based on PLS-DA analysis. For both PLS-DA and PCA analyses count matrices normalized and transformed to variance-stabilized expression levels were used.

The determination coefficient (R2) from partial least squares regression (PLSR) models were used to decipher which clinical parameter was better explained by the grouped top DE sRNAs. Permutation tests (1000 permutations) were applied for statistical significance purposes. Correlation analyses between qPCR-validated sRNAs, NfL, and clinical data were performed using partial correlation test including age and sex as confounding factors.

A 5% of statistical significance level was considered across all analyses.

### qRT-PCR validation

Half of the volume of EVs and NonEVs pooled fractions was used for RNA isolation as described in the section “RNA extraction and small RNA sequencing” with minor modifications. Prior to RNA sample lysis using RPL buffer, synthetic RNA spike-ins (UniSp2, UniSp4, UniSp5 RNA Spike-in mix; Qiagen) were added and used as quality controls for RNA isolation, in accordance with the kit protocol. Two μL of isolated RNA were used for reverse transcription (RT) and cDNA synthesis using the miRCURY LNA RT Kit (Qiagen) according to the manufacturer’s instructions adding the RNA spike-in UniSp6 as quality control for RT. The obtained cDNA was further diluted 1:10 for qPCR and, 3 μL were used as template in a 10 μL final volume reaction using the miRCURY LNA SYBR Green PCR Kit (Qiagen) in a StepOnePlus Real-Time PCR System (Applied Biosystems). For miRNAs validation, specific LNA PCR primer assays (Qiagen) for hsa-miR-451a, hsa-miR-21-5p, hsa-miR-26a-5p, hsa-let-7a-5p, hsa-miR-27a-3p (YP02119305, YP00204230, YP00206023, YP00205727, and YP00206038, respectively) were used. For tRFs validation, custom LNA PCR primers were designed and synthesized by Qiagen for hsa-tRF-Glu-CTC (5’-TCCCTGGTGGTCTAGTGGTTAGGATTCGGCG –3’), hsa-tRF-Gly-GCC (5’-GCATTGGTGGTTCAGTGGTAGAATTCTCGC –3’), and hsa-tRF-Lys-TTT (5’-TAGCTCAGTTGGTAGAGC-3’). For each determination, reactions were performed in triplicate.

Due to the absence of universal endogenous sRNA normalizers in EVs, miR-100-5p was selected as the housekeeping gene since it was one of the most stable sRNAs in terms of least variability across all the sequenced samples and highly expressed. For the analysis of deregulation based on relative quantification (RQ) of qRT-PCR data, linear mixed-effects models (LMM) were applied that accounted for the different sources of variation derived from the experimental design. Age and sex were considered as confounding factors. For the longitudinal study of the selected sRNAs validated, sensitivity analysis was done before the comparison between longitudinal time-points. In this sense, control samples, P-HD mutation carriers at the baseline time-point, and P-HD mutation carriers at the follow-up time-point were run in parallel. Progression of RQ values was evaluated with the LMM models, which accounted as well here for the correlation structure of the longitudinal nature data.

The diagnostic accuracy of validated sRNAs was explored by receiver operating characteristic (ROC) curves. For the analysis of sRNA biosignatures, logistic binary regression was applied, and the obtained estimated probabilities were used to build ROC curves. Area under the ROC curve (AUC) and its 95% confidence interval were used as the diagnostic accuracy metric.

### EV-TRACK

We have submitted all relevant data of our experiments to the EV-TRACK knowledgebase^68^ (EV-TRACK ID: EV230044).

## Acknowledgments

The authors would like to thank all patients at the Hospital de la Santa Creu I Sant Pau and volunteers involved in the study for providing plasma samples. In particular, we want to thank the nurses Claudia Palomera and Andrea García for their involvement in sample collection and processing. The authors would also like to thank Dr. Maria Yañez-Mo (Unidad de Investigación, Hospital Sta Cristina; Departamento Biología Molecular/CBM-SO, UAM) for kindly gifting us antibodies anti-CD9 and anti-CD63 for flow cytometry assays. We wish to thank José Amable Bernabé for his fast measurements of EVs by NTA. It has been performed by the ICTS “NANBIOSIS”, Unit 6, unit of the CIBER in Bioengineering, Biomaterials & Nanomedicne (CIBER-BBN) at the Barcelona Materials Science Institute. Also, we thank Martí de Cabo for acquiring excellent images of EVs by cryo-EM (Servei de Microscòpia, Universitat Autónoma de Barcelona). We thank the staff of Genomics Unit at the CRG for the preparation of sRNA libraries and sRNA sequencing. Finally, we wish to gratefully acknowledge Luisa Mariscal and Sonia Jansa from bioNova for generously providing advice and support regarding sRNA isolation and validation.

## Supplementary data

**Supplementary data Fig. 1:**
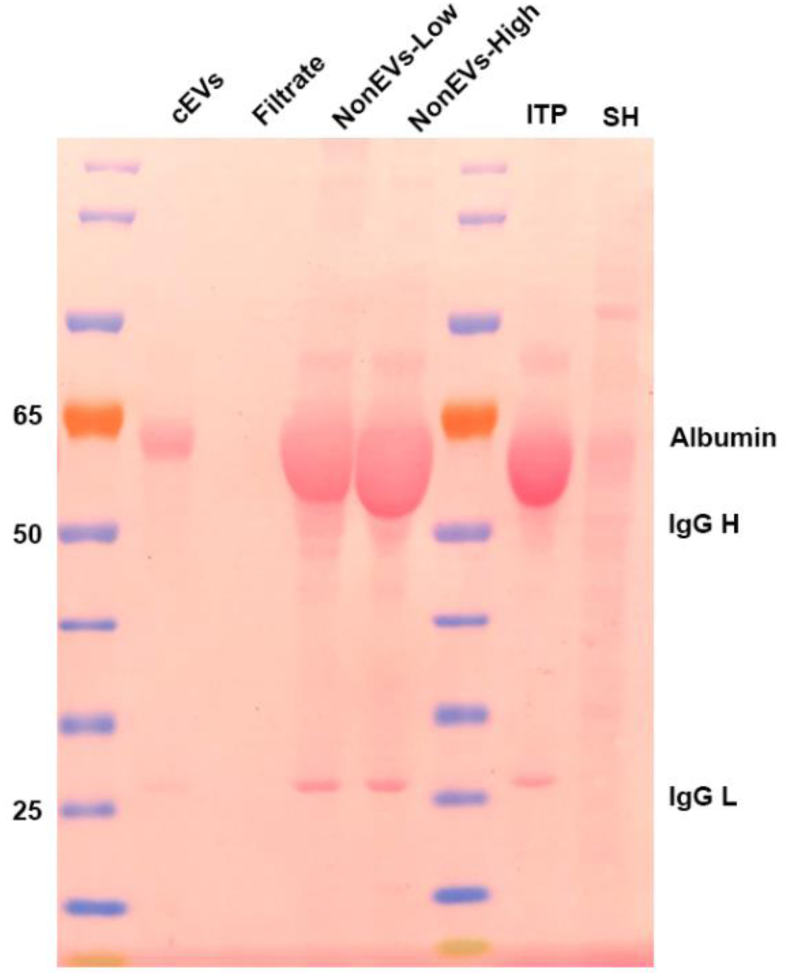
Ponceau S staining of isolated fractions. (A) Albumin, IgG heavy and IgG Light chain were analysed in the transferred membrane. Initial total plasma (ITP) and SH-SY5Y cell lysate (SH) were used as positive and negative controls, respectively.

**Supplementary data Fig. 2:**
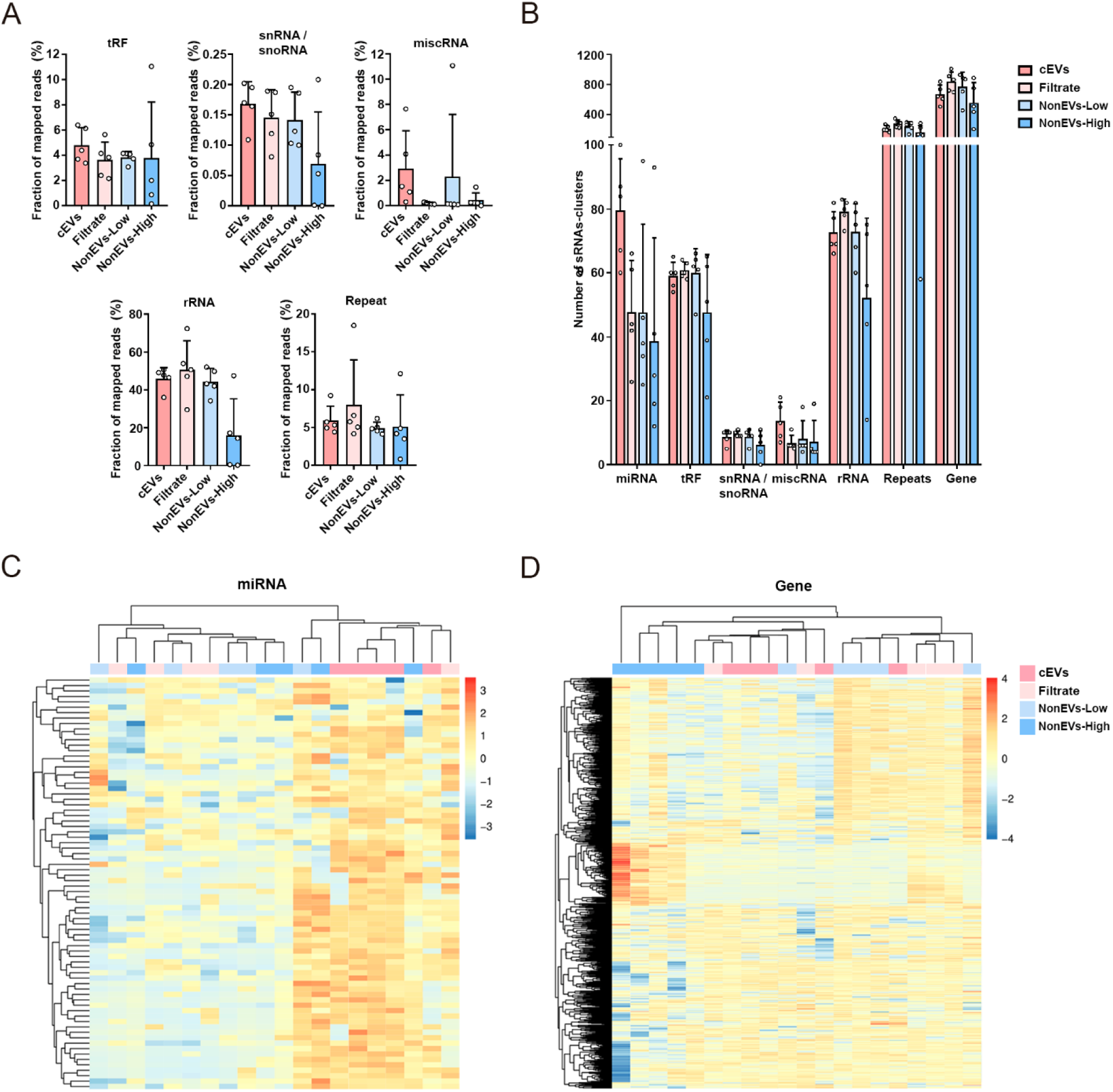
sRNA analysis of plasma subfrations using SeqCluster tool. (A) Fraction of reads mapping onto tRF, snRNA/snoRNA, miscRNA, rRNA and Repeats fragments per plasma subfraction. (B) Absolute number of different types of sRNA-clusters identified per plasma subfraction. Heatmaps of hierarchical clustering analysis of sRNAs mapped onto miRNAs (C) and gene fragments (D). Samples of each subfraction, n=5. Mean ± SD is shown.

**Supplementary data Fig. 3:**
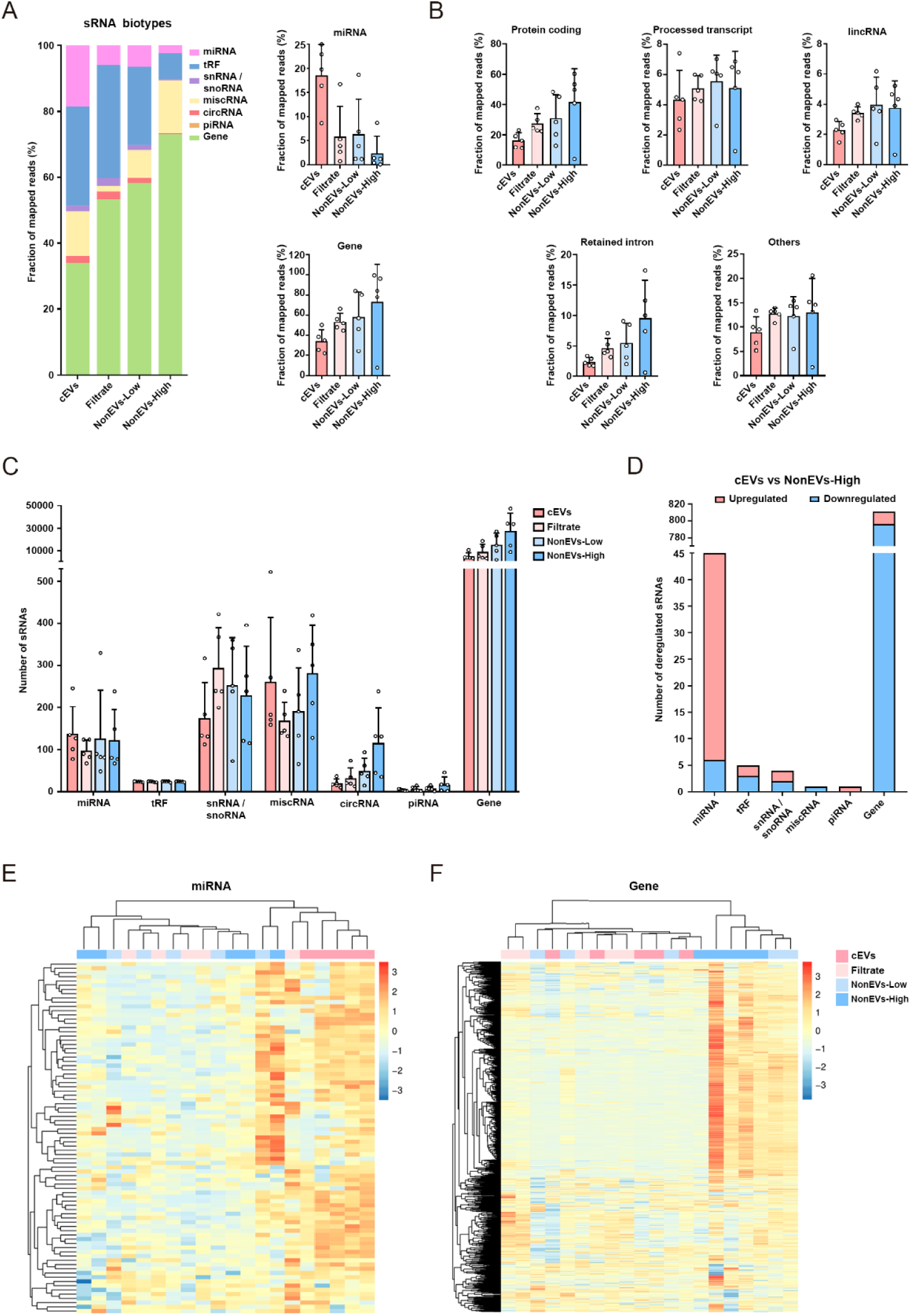
sRNA analysis of plasma subfrations using Excerpt tool. (A) Fraction of reads that align to small RNA types are shown per plasma subfraction. (B) Fraction of reads that align to gene fragments subclassified in Protein coding, Processed transcript, lincRNA, Retained intron fragments, and Others per plasma subfraction. (C) Absolute number of different types of sRNAs identified per plasma subfraction. (D) Total number of sRNAs differentially expressed between cEVs and NonEVs-High subfractions (p-adj <0.05). Heatmaps of hierarchical clustering analysis of sRNAs identified as miRNAs (D) and Gene fragments (E). Samples of each subfraction, n=5. Mean ± SD is shown.

**Supplementary data Fig. 4:**
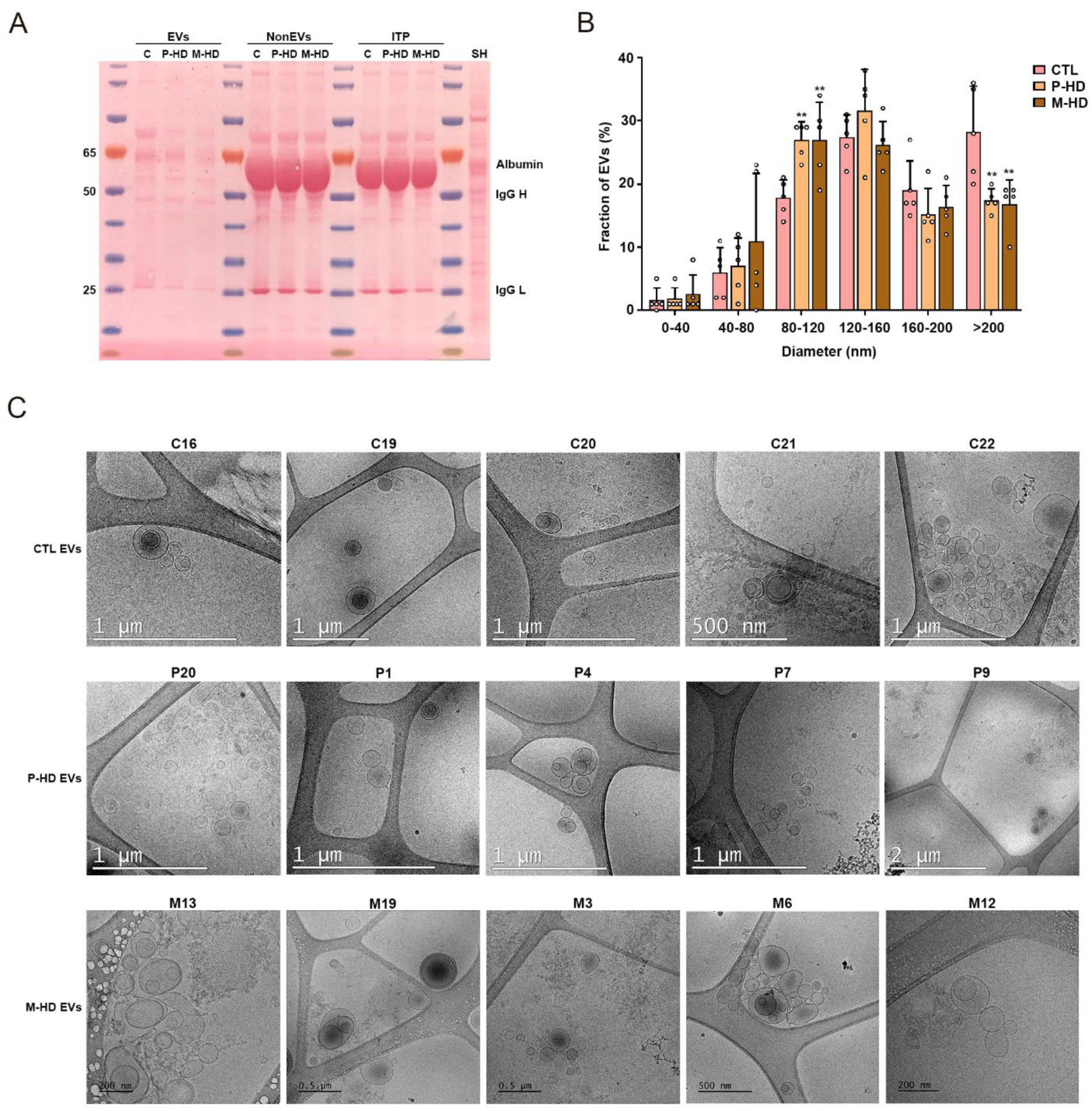
**Characterization of EV fraction and NonEVs fraction isolated from P-HD, M-HD and CTL plasma**. (A) Albumin, IgG heavy and IgG Light chain were analysed by Pounceau Red staining in the representative transferred membrane. Initial total plasma (ITP) and SH-SY5Y cell lysate (SH) were used as positive and negative controls, respectively. (B) Size distribution of EVs diameter by NTA (n=5 per group). All data are represented as Mean ± SD. Significant differences between P-HD and M-HD versus CTL groups using are presented with *** (adj P <0.001), ** (adj P <0.01), and * (adj P <0.05). (C) Examples of images of cryo-EM from the five samples per group (n=5 per group).

**Supplementary data Fig. 5:**
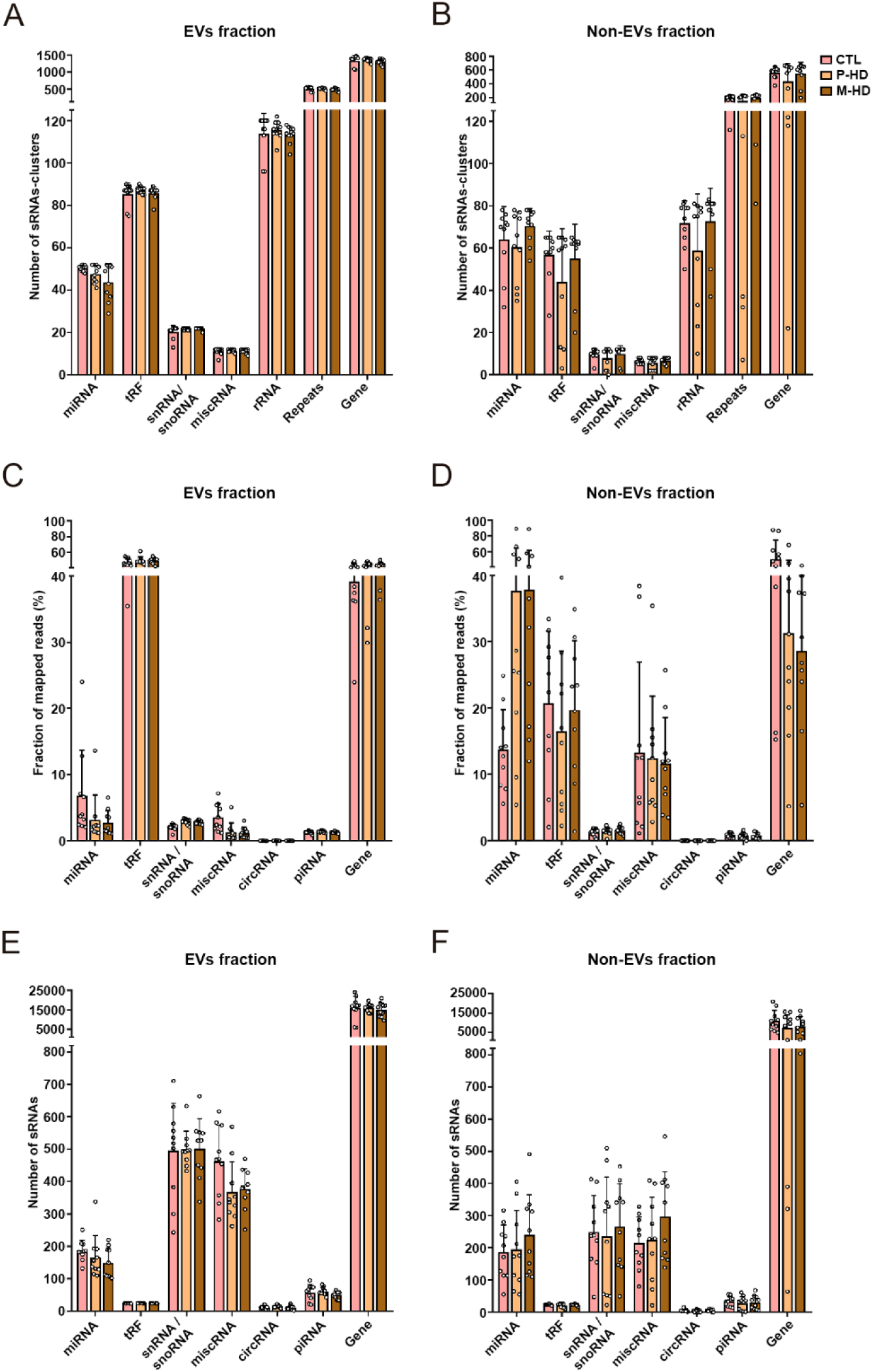
sRNA profiles in EVs and Non-EVs fractions from P-HD, M-HD, and CTL plasma samples. Using SeqCluster tool, absolute number of different types of sRNAs-clusters identified per plasma cohort in (A) EVs fractions and (B) NonEVs fractions. Using ExceRpt tool, fraction of reads that align to small RNA types are shown per plasma cohort in (C) EVs fractions and (D) NonEVs fractions; and absolute number of different types of sRNAs identified per plasma cohort in (E) EVs fractions and (F) NonEVs fractions. Samples per cohort, n=10. Mean ± SD is shown.

**Supplementary data Fig. 6:**
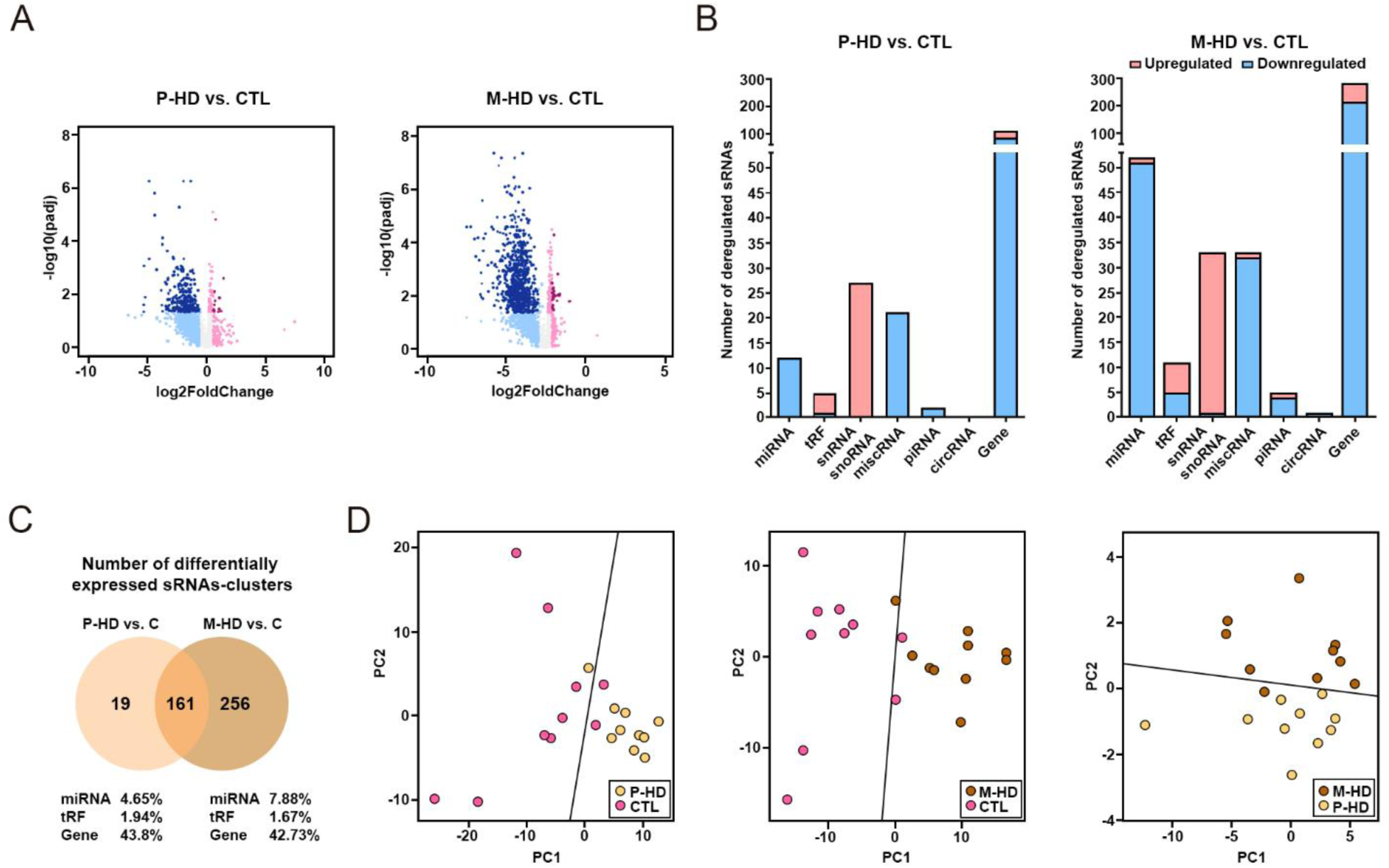
sRNAs from HD patients are deregulated in EVs fractions in comparison to CTL samples, using ExceRpt tool. (A) Volcano plots showing differentially expressed sRNAs in P-HD-EVs versus CTL-EVs fractions and M-HD-EVs versus CTL-EVs fractions. Orange dots represent significantly deregulated genes (|log2FoldChange|>0.58, adjusted P<0.05). (B) Total number of sRNAs differentially expressed between P-HD-EVs versus CTL-EVs fractions and M-HD-EVs versus CTL-EVs fractions. ExceRpt tool was used. (C) Venn Diagram of differentially expressed sRNAs between P-HD vs CTL and M-HD vs CTL, showing the number of overlapped dysregulated sRNAs between both comparisons. (D) PCA plots constructed with top DE sRNAs (with PLS-DA based VIP > 0.8) that most contribute to discriminate between disease conditions and healthy individuals. The solid line is the linear discriminant function that best separates disease conditions. n=10 per cohort, adjusted P<0.05.

**Supplementary data Fig. 7:**
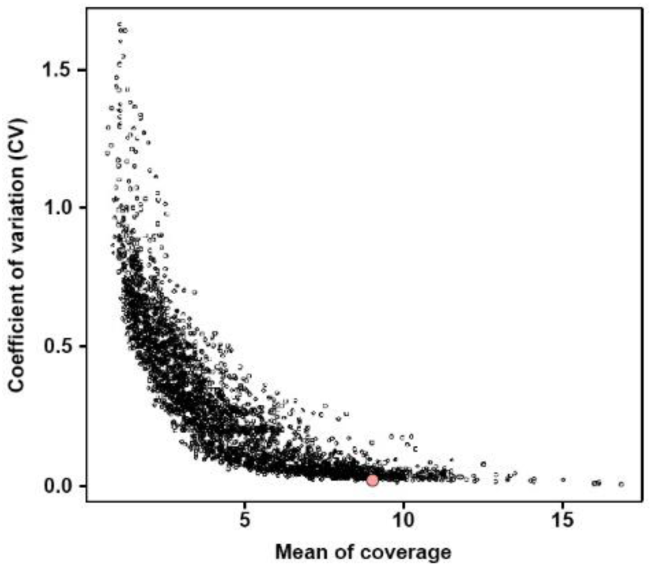
Selection of miR-100-5p as the reference gene for validations by qRT-PCR. Pink dot represents miR-100-5p.

**Supplementary data Fig. 8:**
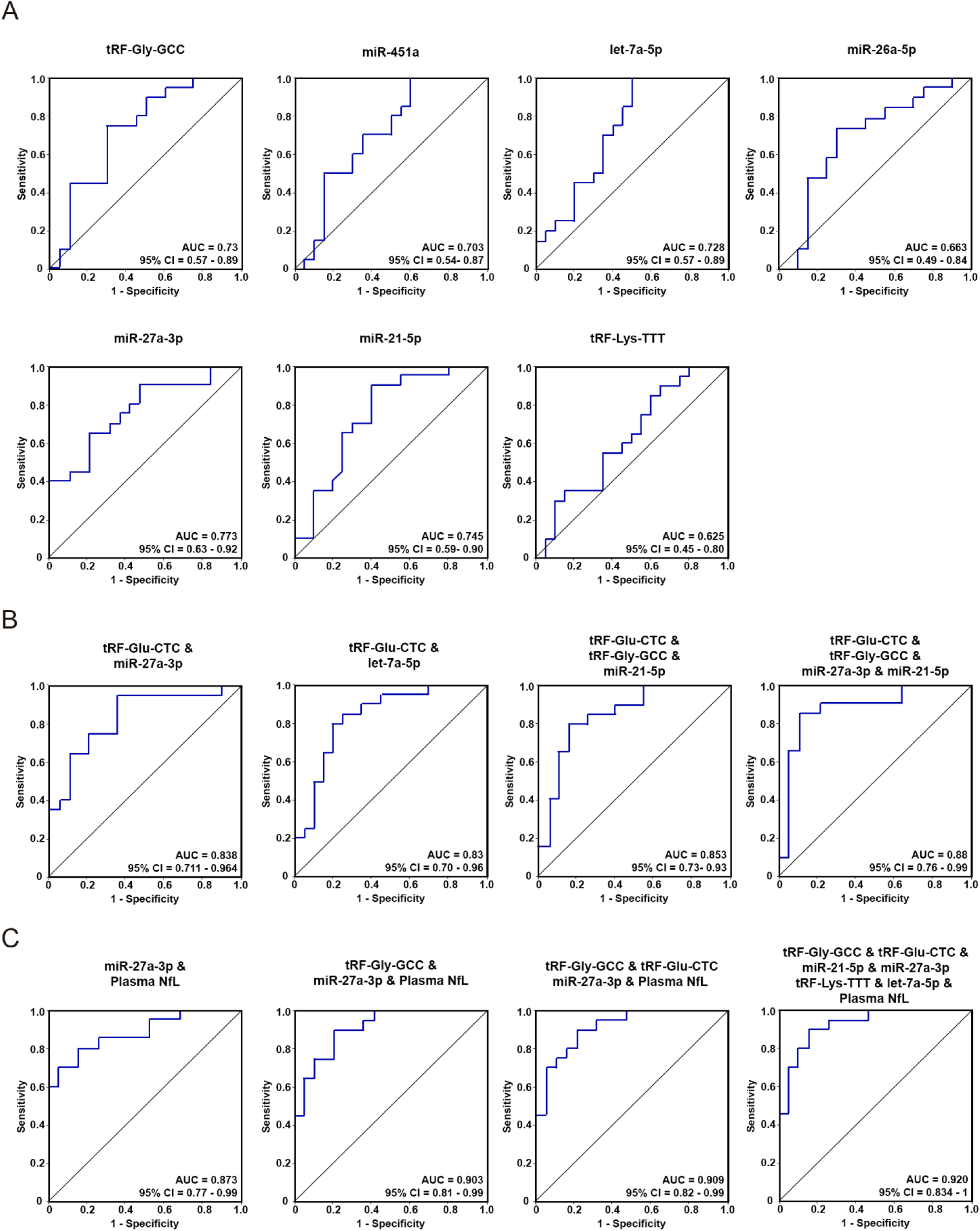
Diagnostic potential analysis of selected sRNAs at premanifest stage. (A) ROC curves analysis of the sensitivity and specificity of individual validated sRNAs between P-HD and CTL samples: tRF-Gly-GCC (p-val=0.013), miR-451a (p-val=0.031), let-7a-5p (p-val=0.014), miR-26a-5p (p-val=0.092), miR-27a-3p (p-val=0.003), miR-21-5p (p-val=0.008), and tRF-Lys-TTT (p-val=0.249). (B) ROC curves analysis of the sensitivity and specificity of novel sRNAs-biosignatures: ensemble of tRF-Glu-CTC and miR-27a-3p (p-val=0.0001); ensemble of tRF-Glu-CTC and let-7a-5p (p-val=0.0001); ensemble of tRF-Glu-CTC, tRF-Gly-GCC and miR-21-5p (p-val=0.0001); and ensemble of tRF-Glu-CTC, tRF-Gly-GCC, miR-27a-3p and miR-21-5p (p-val=0.0001). (C) ROC curves analysis of the sensitivity and specificity of novel sRNAs-biosignatures in combination with plasma NfL: ensemble of miR-27a-3p and plasma NfL (p-val=0.0001); ensemble of tRF-Gly-GCC, miR-27a-3p and plasma NfL (p-val=0.0001); ensemble of tRF-Gly-GCC, tRF-Glu-CTC, miR-27a-3p and plasma NfL (p-val=0.0001); and ensemble of tRF-Gly-GCC, tRF-Glu-CTC, miR-21-5p, miR-27a-3p, tRF-Lys-TTT, let-7a-5p and plasma NfL (p-val=0.0001). n=20 per cohort.

**Supplementary data Fig. 9:**
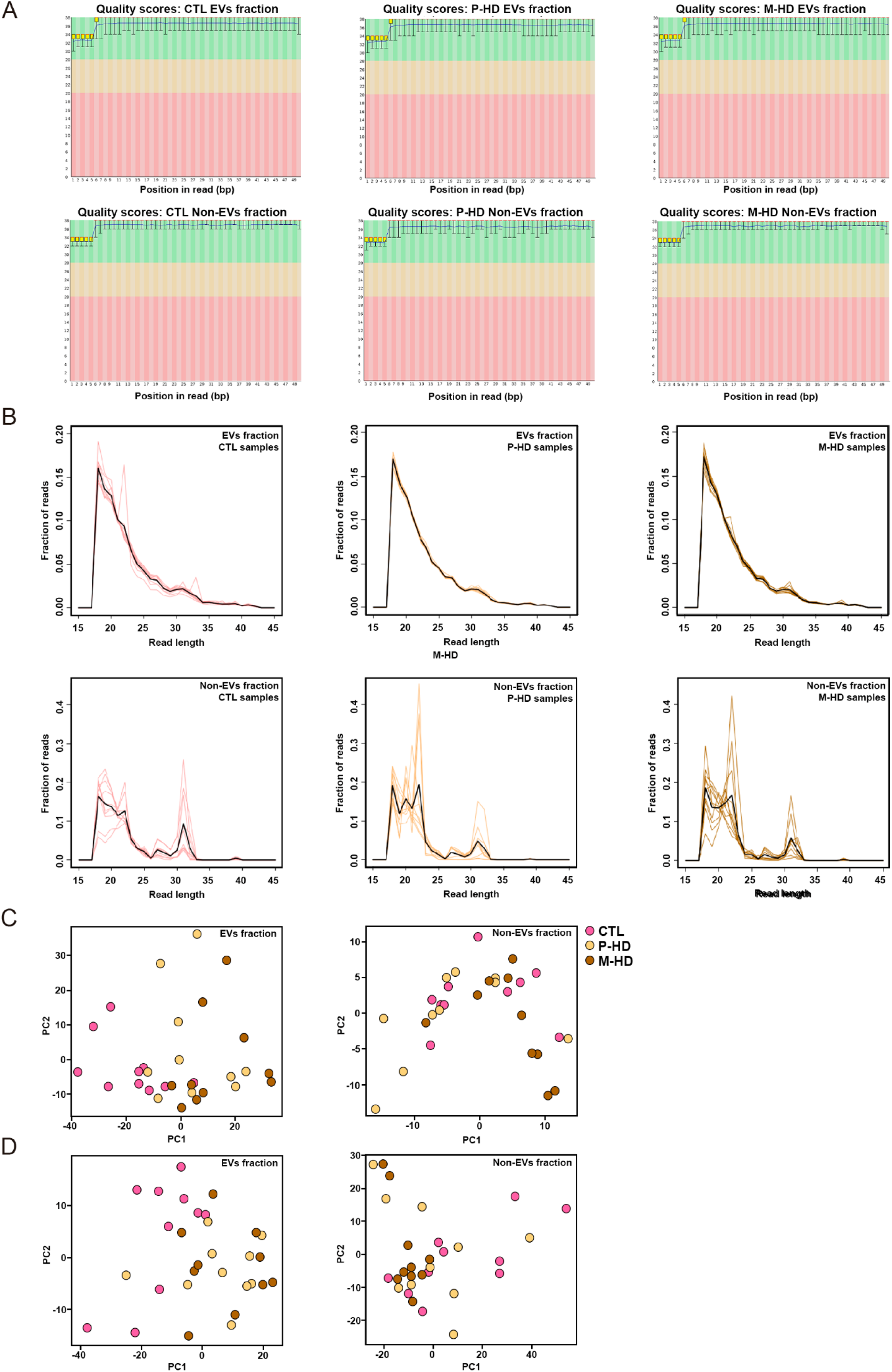
Quality check of RNA sequencing data. (A) Representative images of sequence quality scores across all bases of EVs fraction and Non-EVs fraction per group. (B) Length distribution of sequence reads found with adapters in EVs and Non-EVs fractions per group. Black line represents the average across samples (n=10 per group). First two components of principal component analysis (PCA) of the normalized sRNA sequence count matrix obtained with SeqCluster tool (C) or ExceRpt tool (D). The normalization procedure is based on the regularized log transformation function from DESeq R package.

